# Neuronal heterogeneity of normalization strength in a circuit model

**DOI:** 10.1101/2024.11.22.624903

**Authors:** Deying Song, Douglas Ruff, Marlene Cohen, Chengcheng Huang

**Affiliations:** Joint Program in Neural Computation and Machine Learning, Neuroscience Institute, and Machine Learning Department, Carnegie Mellon University, Pittsburgh, PA; Center for the Neural Basis of Cognition, Pittsburgh, PA; Department of Neurobiology and Neuroscience Institute, University of Chicago, Chicago, IL; Department of Neuroscience and Department of Mathematics, University of Pittsburgh, Pittsburgh, PA

## Abstract

The size of a neuron’s receptive field increases along the visual hierarchy. Neurons in higher-order visual areas integrate information through a canonical computation called normalization, where neurons respond sublinearly to multiple stimuli in the receptive field. Neurons in the visual cortex exhibit highly heterogeneous degrees of normalization. Recent population recordings from visual cortex find that the interactions between neurons, measured by spike count correlations, depend on their normalization strengths. However, the circuit mechanism underlying the heterogeneity of normalization is unclear. In this work, we study normalization in a spiking neuron network model of visual cortex. The model produces a range of neuronal heterogeneity of normalization strength and the heterogeneity is highly correlated with the inhibitory current each neuron receives. Our model reproduces the dependence of spike count correlations on normalization as observed in experimental data, which is explained by the covariance with the inhibitory current. We find that neurons with stronger normalization are more sensitive to contrast differences in images and encode information more efficiently. In addition, networks with more heterogeneity in normalization encode more information about visual stimuli. Together, our model provides a mechanistic explanation of heterogeneous normalization strengths in the visual cortex, and sheds new light on the computational benefits of neuronal heterogeneity.

## Introduction

Understanding how the brain integrates and extracts information from multiple stimuli has long been a central focus in neuroscience. In visual cortex, neurons in the primary visual area respond to local features of stimuli, such as orientation and moving direction, while neurons in the higher-order visual areas have broader receptive fields and extract global features of visual stimuli. The responses of visual neurons to multiple stimuli have been well characterized by a phenomenon known as normalization, where neurons respond sublinearly. Normalization has been observed across brain regions (Heeger, 1992; Louie et al., 2011; Ohshiro et al., 2011), sensory modalities (Olsen et al., 2010; Rabinowitz et al., 2011) and species (Barbera et al., 2022; MacEvoy et al., 2009; Ni et al., 2012), and has been regarded as a canonical computation in nervous system (Carandini and Heeger, 2012). Normalization has also been thought to be the fundamental mechanism through which selective attention acts to enhance neuronal responses to attended stimulus (Reynolds and Heeger, 2009).

It has long been demonstrated that the strength of normalization is variable across neurons (Lee and Maunsell, 2009; Ni et al., 2012; Ruff et al., 2016). Some neurons are suppressed by the addition of a non-preferred stimulus, while some neurons show additive responses to multiple stimuli. Recent population recordings from multiple visual cortical areas of macaque monkeys reveal that the interactions between neurons, measured by spike count correlations, depend on their normalization strengths (Ruff et al., 2016; Verhoef and Maunsell, 2017). Spike count correlations are shaped by network connectivity and dynamical state, and impose strong constraints on circuit models (Hennequin et al., 2018; Huang et al., 2019; Huang, 2021; Ocker et al., 2017). In addition, spike count correlations constrain the amount of information encoded by a neuronal population (Kohn et al., 2016) and can reflect perceptual inference (Bànyai and Orbàn, 2019). Therefore, an understanding of the network mechanisms underlying the relationship between normalization strength and neuronal correlations is likely to provide insights into the neural population code of the integration of multiple stimuli.

Despite the success of the phenomenological models of divisive normalization at reproducing the firing rates of neurons under different stimulus conditions, the neurophysiological basis of normalization remains unclear. Recently, mechanistic circuit models have been developed to account for the sublinear response properties or reproduce the divisive scaling of responses (Heeger and Mackey, 2019; Heeger and Zemlianova, 2020; Lindsay et al., 2019; Rubin et al., 2015). However, they focus on modeling the trial-averaged neuronal responses in homogeneous neural populations and do not consider the trial-to-trial correlations between neurons and the heterogeneity of normalization strength. Recent statistical models of normalization suggest that the strength of normalization impacts the spiking variability of individual neurons and the correlations between neurons, however, they do not take into account network interactions that shape both normalization and neural correlations (Coen-Cagli and Solomon, 2019; Weiss et al., 2023).

In this work, we study normalization in a two-layer network of spiking neurons, modeling the primary visual cortex (V1) and a higher-order visual area (V4 or MT). The network produces internally generated spiking variability due to a balance of strong excitation and inhibition (Huang et al., 2019; Van Vreeswijk and Sompolinsky, 1996). The neurons in our model exhibit a range of normalization strengths in response to multiple stimuli. Our model reproduces the dependence of spike count correlations on normalization strength as observed in experimental data (Ruff et al., 2016). Interestingly, we identify the inhibitory current to be the major determinant of normalization strength and its relationship with spike count correlations. Further, we demonstrate that neurons with stronger normalization are more sensitive to contrast differences of visual stimuli and encode information more efficiently. Networks with more heterogeneity in normalization encode higher information about visual stimuli, demonstrating the computational benefits of neuronal heterogeneity.

## Results

We use a two-layer spiking neuron network model, with the feedforward layer modeling V1 neurons and the recurrent layer modeling V4 or MT area (Figure 1A). Two Gabor images of orthogonal orientations are presented to the network. The V1 neurons are modeled as linear-nonlinear-Poisson neurons with Gabor receptive fields oriented at their preferred orientation, determined from a superimposed pinwheel orientation map (Kaschube et al., 2010). There are two populations of V1 neurons, V1_1_ and V1_2_, each of which has a non-overlapping receptive field centering on each Gabor image, respectively.

**Figure 1.**
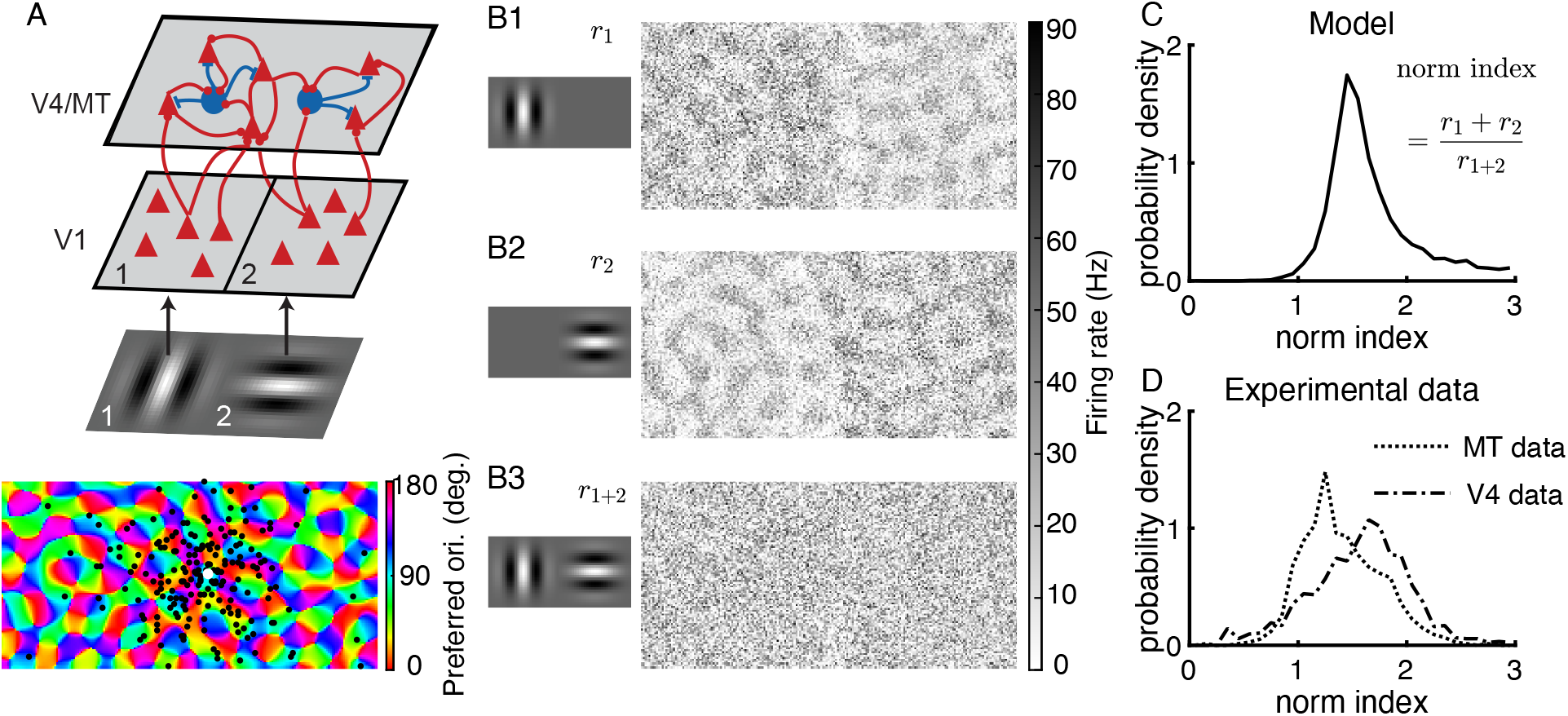
Model schematic, activation patterns, and distributions of normalization indices. (**A**) Model schematic. V4/MT area is modeled as a spatially ordered spiking neuron network of excitatory and inhibitory neurons. V1 neurons are modeled as linear-nonlinear-Poisson neurons with Gabor receptive fields. The two sub-populations of V1 neurons, V1_1_ and V1_2_, have non-overlapping receptive fields, each centering on one Gabor image. The preferred orientations of both V4/MT and V1 neurons are assigned according to pinwheel orientation maps (bottom panel). Bottom: Locations of postsynaptic excitatory neurons (black dots) of one example presynaptic excitatory neuron (white dot) in the V4/MT layer. The majority of the connections are local in space and a small portion of the connections are between similarly tuned excitatory neurons across the whole network (see Methods). (**B**) Firing rate patterns of the model V4/MT neurons when one Gabor image is presented at location 1 (**B1**) or location 2 (**B2**), or when both images of orthogonal orientations are presented together (**B3**). (**C**) The model V4/MT neurons exhibit a wide range of normalization indices. The normalization index is defined as the sum of the neuron’s firing rates in response to each image presented individually (**B1, B2**), divided by its firing rate when both images are presented together (**B3**). (**D**) The normalization indices of neurons recorded from macaque V4 and MT areas (data from Ruff et al. (2016)).

The V4/MT layer is a recurrent network of excitatory and inhibitory neurons, each modeled as an exponential integrate-and-fire neuron. Neurons from the V4/MT layer receive inputs from both V1 populations and respond to both images (Figure 1B). There are two types of connections in both feedforward projections from V1 to V4/MT and and recurrent projections within the V4/MT layer (Figure 1A), following anatomical findings from visual cortex (Angelucci et al., 2002; Bosking et al., 1997; Malach et al., 1993; Mariño et al., 2005). The majority of connections are local, of which the connection probability decays with distance. We choose the spatial scales of excitatory and inhibitory projections to be the same, consistent with anatomy (Mariño et al., 2005). A small portion of connections are long-range, meaning that their connection probability does not depend on distance, but they connect between similarly tuned excitatory neurons. The spatially dependent connections allow the V4/MT neurons to retain location information of the two images, while the tuning specific long-range connections increase the tuning selectivity and the size of receptive fields of V4/MT neurons.

The V4/MT network admits a stable and asynchronous solution with homogeneous V1 inputs, similar to the classic balanced network model (Renart et al., 2010; Van Vreeswijk and Sompolinsky, 1996) (Supp Fig S1). The balance between strong excitation and inhibition generates Poisson-like spiking variability in individual neurons. The distribution of firing rates is lognormal due to the expansive transfer function of neurons with large input fluctuations (Roxin et al., 2011). With orientation-tuned inputs from V1, the V4/MT model neurons capture several features of neuronal responses in visual cortex, as shown in our previous work (Huang et al., 2022). The tuning curves are heterogeneous with various widths and magnitudes (Ringach et al., 2002). The spike count correlations are higher between similarly tuned neurons (Cohen and Maunsell, 2009; Gu et al., 2011), and depend on stimulus orientation (Hennequin et al., 2018; Lin et al., 2015; Ponce-Alvarez et al., 2013).

### Broad distribution of normalization strengths in the model and data

We analyze the network responses to two Gabor images with orthogonal orientations, which are known to evoke the normalization mechanism in the visual cortex (Busse et al., 2009; Ni et al., 2012; Ruff et al., 2016; Verhoef and Maunsell, 2017). Neurons’ responses to two stimuli presented together tend to be much less than the linear sum of the responses to each stimulus when presented individually. This phenomenon has been observed across the visual hierarchy in macaque monkeys (Ruff et al., 2016) as well as in the primary visual cortex in cat (Busse et al., 2009) and tree shrew (MacEvoy et al., 2009).

When presented with one image, neurons that prefer the orientation of the image are activated across the V4/MT network, resulting in local patches of active regions following the pinwheel map of orientation preference (Figure 1B1, B2). Neurons in the same spatial location of the presented image respond with higher rates since they receive more local feedforward inputs. When both images are presented together, there is no clear spatial structure of population activation pattern of firing rates (Figure 1 B3).

We define normalization index of each neuron as the sum of the neuron’s firing rates in response to each image presented individually, divided by its firing rate when both images are presented together (Eq. 14), following the definition from our previous experimental work (Ruff et al., 2016). A normalization index of one means that the neuron’s response to multiple stimuli is a linear summation and a normalization index larger than one means a sublinear summation. A larger normalization index means stronger normalization.

Our model V4/MT neurons exhibit a wide range of normalization indices with the majority being between one and two (Figure 1C). The normalization indices of our model neurons span a similar range as those of neurons recorded from macaque V4 and MT areas (Figure 1D; Ruff et al. (2016)). The normalization index of a neuron is independent from its tuning preference in both our model and data (Figure S2). The wide distribution of normalization indices is a robust phenomenon in networks with strong recurrent connections (Figure S3). Even in random networks with no spatial or tuning dependent connections, there is a spread of normalization indices due to random connections when two input populations project to the whole network (Figure S3A,B magenta). The distribution widens when the two input populations project to distinct sets of target neurons (Figure S3B cyan). In our model, the two images of orthogonal orientations activate different sets of V4/MT neurons, which is similar to the case with small overlap of input projections in the random networks (Figure S3B). In contrast, networks with weak recurrent connections produce narrow distributions of normalization strengths (Figure S4). Being able to reproduce the range of neuronal heterogeneity of normalization in our model, we next examine how neurons with different normalization strengths interact with each other.

### Spike count correlations between neurons depend on their similarity of normalization strength

The neuronal heterogeneity of normalization strength in our model is shaped by network connectivity and neuronal transfer functions. Both factors are known to determine the interactions between neurons (Huang et al., 2019; Ocker et al., 2017). Next we examine how correlations between neurons depend on neurons’ normalization indices. We use spike count correlations to measure the interactions between neurons, which is commonly used to measure the trial-to-trial co-fluctuations of neuronal responses in experiments (Cohen and Kohn (2011); see Methods).

We find that spike count correlations between neurons depend on their normalization indices in the model in a way that is qualitatively consistent with the neuronal correlations measured in visual cortex (Figure 2). First, neurons with similar normalization indices have higher spike count correlations than those with different normalization indices (diagonal vs. off-diagonal elements in Figure 2A1, B1 and C1). For neuron pairs of the same average normalization indices, their spike count correlations decrease as the difference between their normalization indices becomes larger (Figure 2A2, B2 and C2). Second, for neuron pairs with similar normalization indices (diagonal elements in Figure 2A1, B1 and C1), their spike count correlations first decrease with their average normalization indices and can increase at large normalization indices in the model and the V4 data (Figure 2A3, B3 and C3). The same pattern of spike count correlations is also observed in neurons recorded from Macaque V1 area (Supp Fig S5). The model also produces the same relationship between spike count correlation and normalization strengths using superimposed Gabor images in addition to two separately presented images (Supp Fig S6). The close match of the dependence of correlations on normalization indices in our model with that in data suggests common circuit mechanisms that determine the heterogeneity of normalization. In contrast, the dependence of neuronal correlations on normalization index is very weak in networks with weak recurrent connections, emphasizing the importance of recurrent connections in determining the strength of normalization (Figure S4).

**Figure 2.**
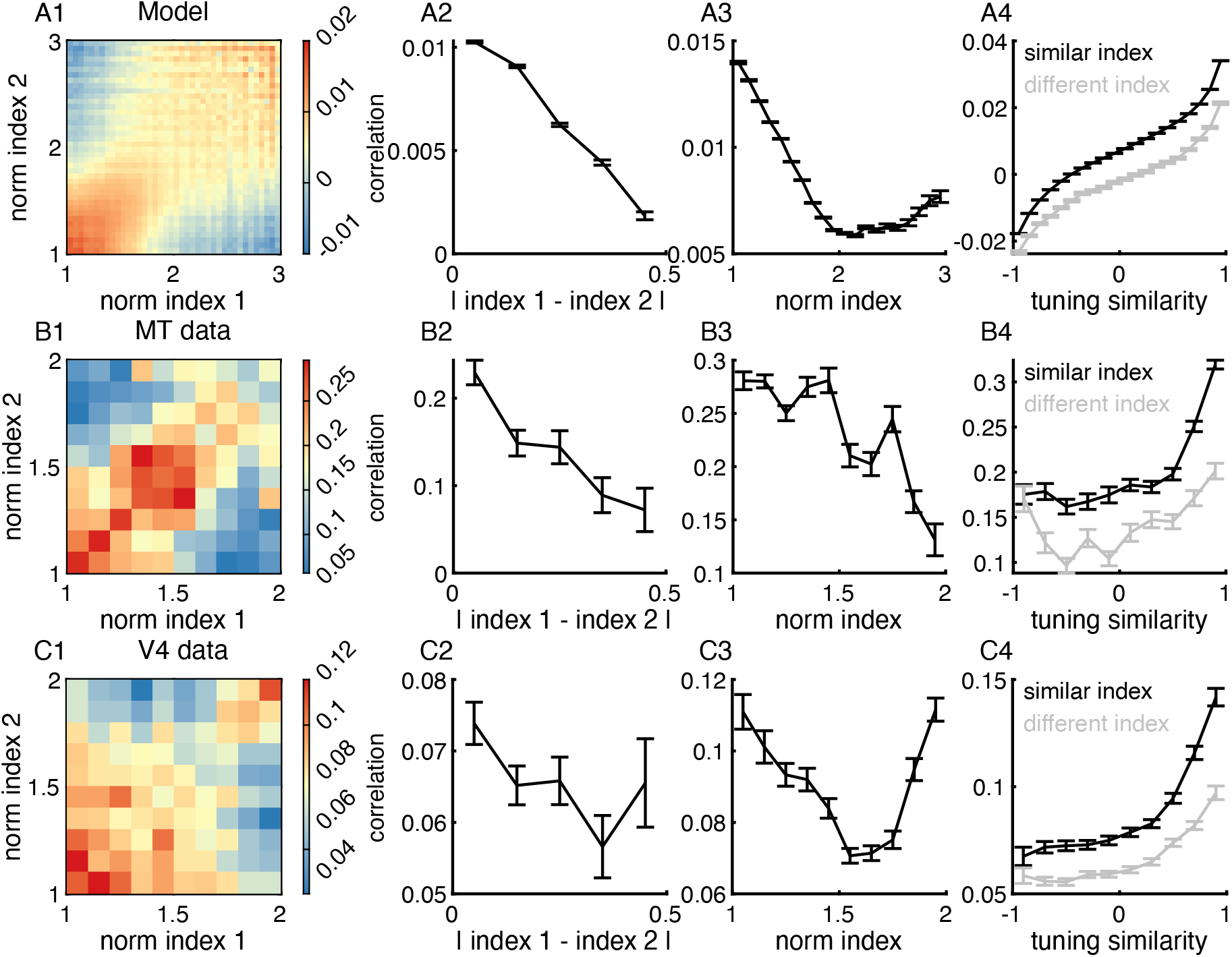
Neurons with similar normalization indices have higher spike count correlations than those with different normalization indices. (**A**) Spike count correlations between a pair of model MT/V4 neurons depend on their normalization indices. (**A1**) Spike count correlations between a pair of neurons as a function of the normalization indices of the pair. (**A2**) For neuron pairs of the same average normalization indices (equal to 1.5), their spike count correlations decrease with the difference in their normalization indices. (**A3**) For neuron pairs of similar normalization indices (difference < 0.5), their spike count correlations decrease with their average normalization indices. (**A4**) Across all levels of tuning similarity, the spike count correlations between neurons with similar normalization indices are consistently larger than those of neurons with distinct normalization indices. (**B**) Same as (A) for neuronal data recorded from the MT area (28 sessions). (**C**) Same as (A) for neuronal data recorded from the V4 area (21 sessions). Error bars represent the SEM. Data in panels B and C are from Ruff et al. (2016).

A potential explanation for the observed dependence of correlations on the similarity of normalization indices is that neurons with similar normalization indices may have more similar tuning preferences than those with different normalization indices. Several previous experimental studies have consistently demonstrated that neurons with similar tuning preferences tend to have higher spike count correlations (Cohen and Maunsell, 2009; Gu et al., 2011; Hennequin et al., 2018; Lin et al., 2015). Therefore, tuning similarity between neurons is a potential confound that underlies the dependence of correlations on normalization indices. However, this is not the case in our model and in data. First, we find no significant correlation between the tuning preference and the normalization index of a neuron in both model and data (Supp Fig S2). Second, we compare the spike count correlations of neuron pairs with either similar or different normalization indices across different magnitudes of tuning similarity, measured as the correlation between the tuning curves of two neurons (Eq. 16). We observe that across all levels of tuning similarity, the spike count correlations between model neurons with similar normalization indices are consistently larger than those of model neurons with distinct normalization indices (Figure 2, A4). We re-analyzed our experimental data and found consistent patterns in neural recordings from all three visual areas, MT (Figure 2 B4), V4 (Figure 2 C4) and V1 (Supp Fig S5).

### Recurrent inhibition best explains the heterogeneity of normalization

Having demonstrated that our model successfully reproduces the distribution of normalization strength and the dependence of spike count correlations on normalization indices observed in visual cortex, we next examine the circuit mechanisms in our model that underlie the neuronal heterogeneity of normalization. We decompose the total current each V4/MT neuron receives into three components: feedforward excitation from V1 neurons, recurrent excitation from other V4/MT excitatory neurons and recurrent inhibition from other V4/MT inhibitory neurons. We find that the normalization index of each V4/MT neuron is strongly and negatively correlated with the normalization index of the inhibitory current, defined in the same way as the normalization index of firing rate (Eq. 15; Figure 3C). This means that neurons with stronger normalization receives relatively more inhibition when both images are presented compared to neurons with weaker normalization. In contrast, the correlations between the normalization indices of the firing rates and those of the feedforward and recurrent excitatory currents, respectively, are much weaker (Figure 3A,B). In particular, the weak correlation between normalization and feedforward excitation indicates that the normalization strength of each neuron in the model is mainly determined by recurrent connections. In addition, there is also strong correlation between the firing rate normalization indices and the average inhibitory current a neuron receives when two images are presented, and only weak correlations with the excitatory currents (Supp Fig S7A). The correlation between normalization and the number of excitatory or inhibitory input connections is also weak (Supp Fig S7B).

**Figure 3.**
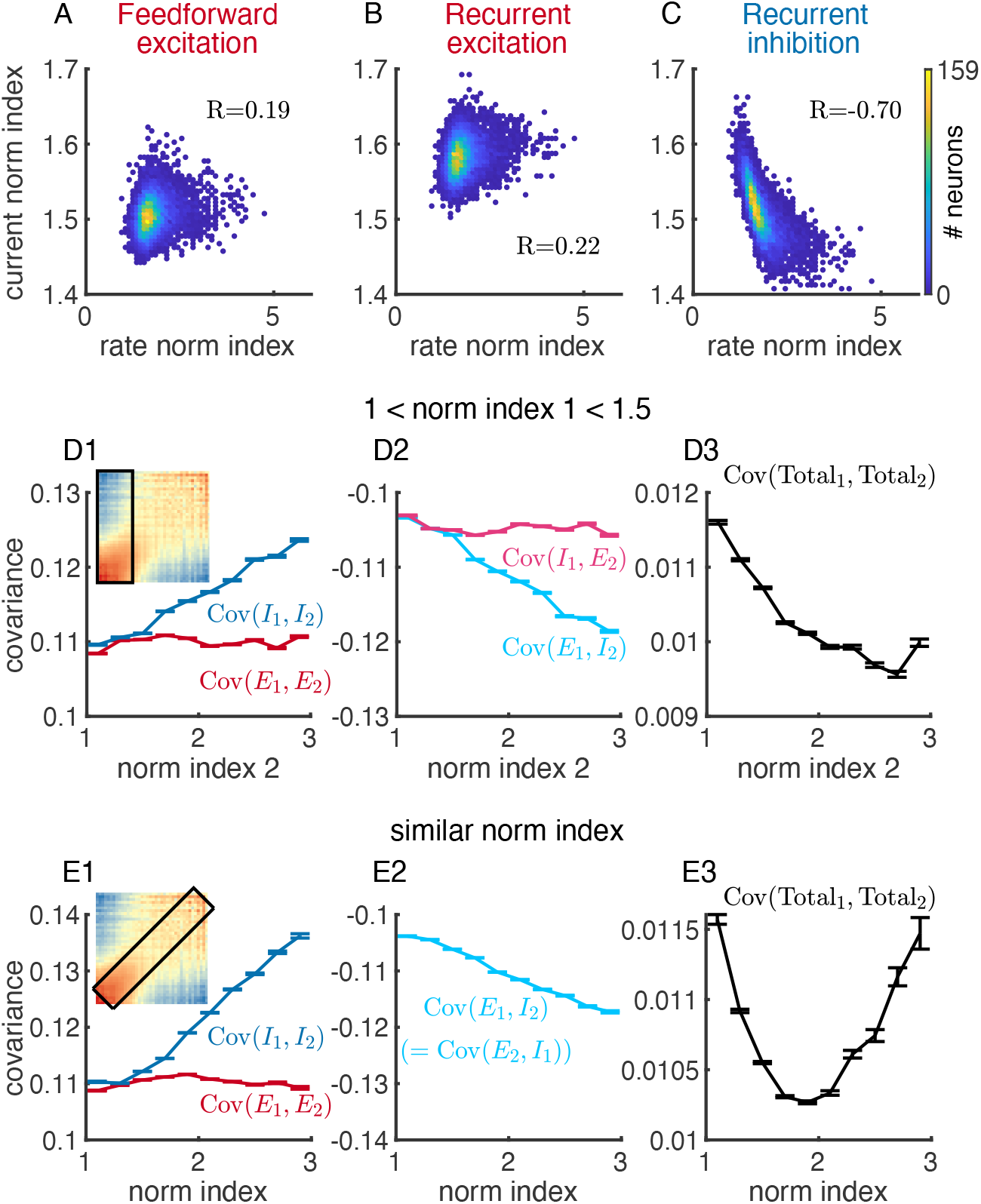
Recurrent inhibition best explains the heterogeneity of normalization. **(A)** The relationship between the normalization indices of neurons’ firing rates and those of the feedforward excitatory currents neurons receive in the model. (**B-C**) Similar to (**A**), but for recurrent excitatory current **(B)** and recurrent inhibitory current (**C**). The normalization index of recurrent inhibitory current is strongly correlated with the firing rate normalization index. (**D**) When restricting the normalization index of neuron 1 in a pair to be between 1 and 1.5, while allowing the normalization index of the neuron 2 to vary, only the covariance components with the inhibitory currents depend on the normalization index of neuron 2 (Cov(*I*_1_,*I*_2_) in **D1** and Cov(*E*_1_,*I*_2_) in **D2**). Here the excitatory current includes both feedforward and recurrent excitation. The covariance between total currents to the pair of neurons, Cov(Total_1_,Total_2_), decreases with the normalization index of neuron 2 (**D3**), consistent with the changes in spike count correlations (D1 inset). Inset in D1: Spike count correlations between a pair of neurons as a function of their normalization indices, same as in Figure 2A1. Black box indicates the range of normalization indices of neuron pairs analyzed in panels **D1-3**. (**E**) Similar to (**D**), but for neuron pairs with similar normalization indices (difference < 0.6), indicated by the black box in E1 inset. Note that Cov(*I*_1_, *E*_2_) is the same as Cov(*E*_1_, *I*_2_) in panel **E2**, as both normalization indices of neuron 1 and neuron 2 vary together.

To investigate if the inhibitory currents also contribute to the dependence of spike count correlations on normalization indices (Figure 2A), we compute the covariance between the excitatory and inhibitory currents a pair of neurons receive. Let Total_*i*_ be the total current neuron *i* receives (*i* = 1, 2), then Total_*i*_ = *E*_*i*_ + *I*_*i*_, where *E*_*i*_ and *I*_*i*_ are the excitatory and inhibitory current, respectively, that neuron *i* receives. Here we combine both feedforward and recurrent excitation in the excitatory current, *E*, since the following covariance analysis yields similar patterns for both current types. The covariance between the total currents of neuron 1 and neuron 2 can then be decomposed into four components

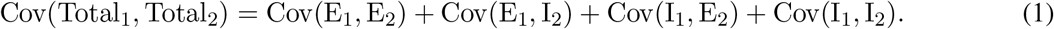

Previous theoretical analysis has shown that in balanced network models both the excitatory and inhibitory inputs to a pair of neurons (Cov(*E*_1_, *E*_2_) and Cov(*I*_1_, *I*_2_), respectively) are correlated which are cancelled by the large negative correlation between the excitatory and inhibitory inputs (Cov(*E*_1_, *I*_2_) and Cov(*I*_1_, *E*_2_)) (Renart et al., 2010). In this way, the total current covariance remains small and neurons in the balanced networks are asynchronous.

We find that only the covariance components with the inhibitory currents depend on normalization indices in our model. We focus our analysis on two cases. First, we restrict the normalization index of neuron 1 in a pair to be between 1 and 1.5, while allowing the normalization index of the neuron 2 to vary (Figure 3D1 inset, black box region). As the normalization index of neuron 2 becomes larger and thus more distinct from neuron 1, the inhibitory current of neuron 2 becomes more correlated with the inhibitory current of neuron 1 (Cov(*I*_1_, *I*_2_) in Figure 3D1) and more negatively correlated with the excitatory current of neuron 1 (Cov(*E*_1_, *I*_2_) in Figure 3D2). In contrast, the covariance components with the excitatory current of neuron 2 is independent of its normalization index (Cov(*E*_1_, *E*_2_) in Figure 3D1 and Cov(*I*_1_, *E*_2_) in Figure 3D2). The covariance, Cov(*E*_1_, *I*_2_), becomes more negative than the increase in Cov(*I*_1_, *I*_2_), resulting in a reduction in the total current covariance, Cov(Total_1_, Total_2_), as the normalization index of neuron 2 increases (Figure 3D3). This is consistent with the changes in spike count correlations (Figure 2A and replicated as the inset of Figure 3D1).

Second, we conduct the same analysis of current covariance components for neuron pairs with similar normalization indices (Figure 3E1, inset, black box region). We observe a similar pattern: the covariance components with the inhibitory currents, Cov(*I*_1_, *I*_2_), Cov(*E*_1_, *I*_2_) and Cov(*I*_1_, *E*_2_), increase in magnitude with normalization index, while the covariance between excitatory currents, Cov(*E*_1_, *E*_2_), is independent of normalization index (Figure 3E1, E2). Note that Cov(*I*_1_, *E*_2_) is the same as Cov(*E*_1_, *I*_2_) in this case as both normalization indices of neuron 1 and neuron 2 vary together. The total current covariance decreases initially as the normalization indices of both neurons increase, and turns to increase when normalization is strong due to the large increase in the covariance between inhibitory currents, Cov(*I*_1_, *I*_2_) (Figure 3E3), consistent with the changes in spike count correlations (Figure 2A1,A3).

In sum, we find that neurons with stronger normalization receives more recurrent inhibition from the network. Their inhibitory currents tend be more correlated with the input currents of other neurons, which allows for better cancellation of current correlations and leads to lower spike count correlations with other neurons.

### Neurons with stronger normalization are more sensitive to contrast differences of images

Our results so far have focused on the population response properties of V4/MT neurons to two images of equal contrast. We next examine the dependence of V4/MT neuron responses on the contrast of each image. We observe that neurons exhibit diverse response tuning to different contrast combinations of the two images (Figure 4, A1-A4). We group neurons by their normalization indices (Eq. 14) and their response selectivity to the two images (Eq. 17). Because of symmetry, here we only present results of neurons that prefer stimulus 1. Therefore, neurons with strong selectivity respond more strongly to stimulus 1, while neurons with weak selectivity respond similarly to both images.

**Figure 4.**
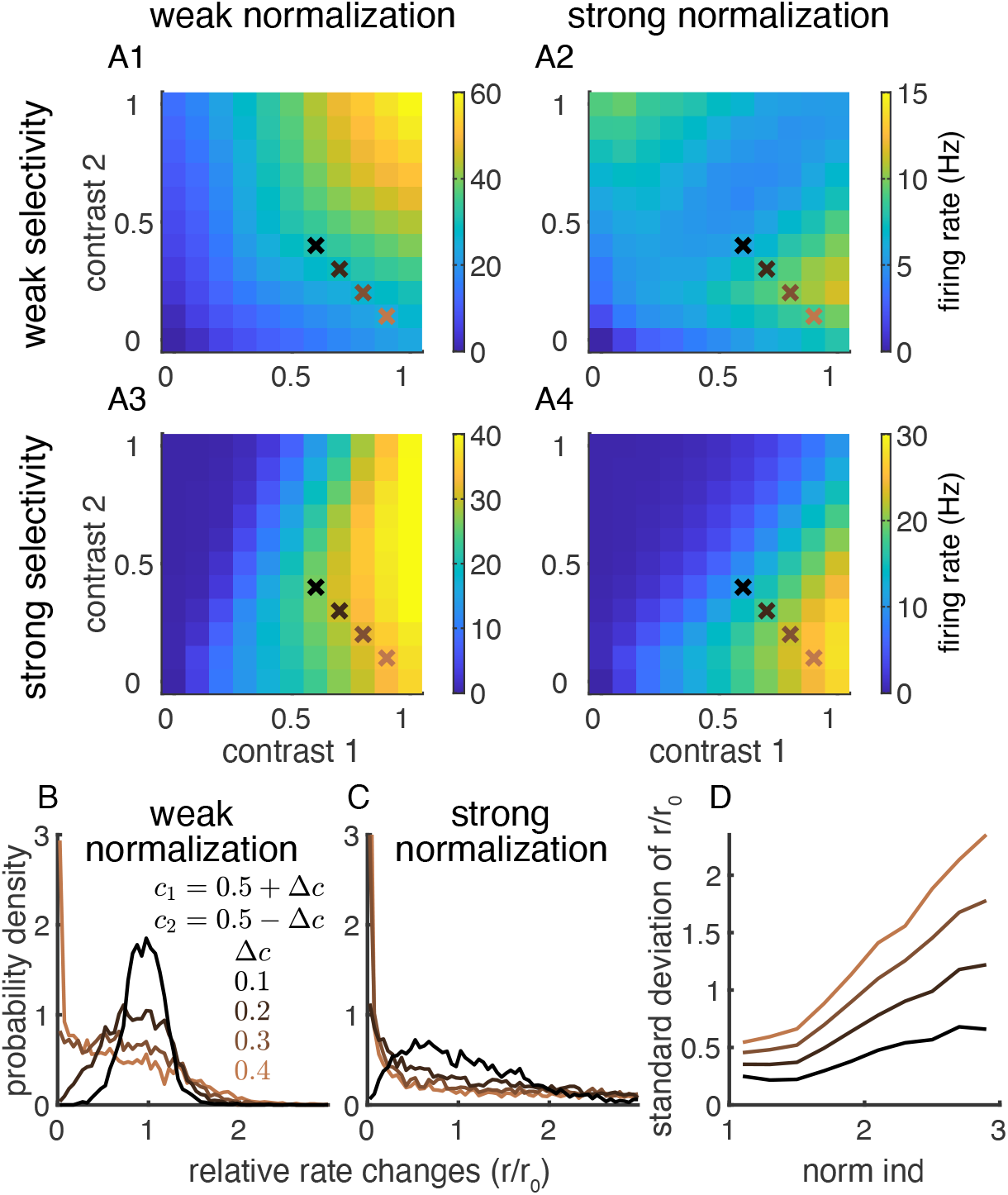
Neurons with stronger normalization are more sensitive to contrast differences of images. (**A**) Neurons exhibit diverse response tuning to different contrast combinations of two Gabor images with orthogonal orientations. (**A1**) The firing rates of neurons with weak normalization and weak selectivity. (**A2**) The firing rates of neurons with strong normalization and weak selectivity. (**A3**) The firing rates of neurons with weak normalization and strong selectivity. (**A4**) The firing rates of neurons with strong normalization and strong selectivity. Neurons that prefer stimulus 1 were selected for the results in A1-4. Crosses indicate the contrast combinations analyzed in B-D with the same color labels. (**B-C**) As contrast difference between the two images increases, the distribution of relative rate changes (*r/r*_0_) of neurons becomes broader. The relative rate changes are neurons’ firing rates to images of a given contrast difference, Δ*c*, divided by their firing rates to images of equal contrast (i.e. Δ*c* = 0). Neurons with strong normalization (**C**) have a much broader distribution of relative rate changes compared to neurons with weak normalization (**B**). (**D**) The standard deviation of rate changes increases with normalization index.

The firing rates of neurons with weak normalization and weak selectivity increase linearly as the contrast of either one of the images increases (Figure 4A1). Thus they respond maximally when both images have high contrast. On the contrary, neurons with strong normalization and weak selectivity are largely suppressed when both images have high contrast, and respond maximally when there is only one image present (Figure 4A2). Neurons with strong selectivity preferentially respond when their preferred image has high contrast (contrast 1), as expected (Figure 4A3-A4). However, neurons with strong normalization are much more suppressed by increasing the contrast of their non-preferred image (contrast 2), and respond maximally when only their preferred image is present (Figure 4A4). Together, neurons with heterogeneous normalization strength and selectivity preferentially respond to different contrast combinations of the two images.

We notice in the contrast dependence of responses that neurons with strong normalization are mostly active when the contrast difference between the two images is large (Figure 4A, lower right corners). To quantify this, we consider conditions where the average contrast of the two images is 0.5 and we gradually increase the difference between the contrasts of the two images. Specifically, we choose contrast 1 as *c*_1_ = 0.5 + Δ*c* and contrast 2 as *c*_2_ = 0.5 − Δ*c*, where 2Δ*c* is the contrast difference between the images (crosses in Figure 4A1-A4). We quantify the relative change in firing rate, *r/r*_0_, by normalizing the firing rate of each neuron with its rate (*r*_0_) when both images have equal contrasts, i.e. Δ*c* = 0. As Δ*c* increases, the distribution of relative rate changes of neurons becomes broader, suggesting that some neurons become more active (*r/r*_0_ > 1) while other neurons become more suppressed (*r/r*_0_ < 1) (Figure 4B,C). Here neurons of both stimulus preferences are included. For the same contrast difference, Δ*c*, neurons with stronger normalization have a much broader distribution of relative rate changes compared to neurons with weaker normalization, suggesting that they are more sensitive to contrast difference (Figure 4B,C). Indeed, the standard deviation of the distributions of relative rate changes increases with the normalization index for each Δ*c* (Figure 4D). This indicates that neurons with strong normalization exhibit much larger changes in their firing rates when there is a contrast difference in the two images.

### Neurons with stronger normalization encode information more efficiently

Normalization has been hypothesized to be computational advantageous because it adapts neurons’ dynamical range of responses and can increase neurons’ sensitivity to changes in stimulus. To quantify the computational benefits of normalization in our model, we measure the linear Fisher information of stimulus parameters from the activity of neurons with different normalization indices. The linear Fisher information measures the accuracy of estimating a stimulus parameter, such as contrast or orientation, from neuron population activity using an optimal linear decoder (Beck et al., 2011; Kohn et al., 2016; Serie`s et al., 2004). The linear Fisher information of a stimulus parameter, *s*, is defined as

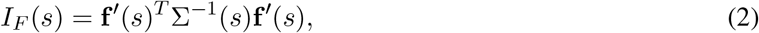

where **f** is the tuning curve function of the neuron population with respect to *s*, ^*′*^ denotes differentiation with respect to *s*, and Σ is the covariance matrix of the population responses. Higher Fisher information means lower threshold in detecting changes in the stimulus parameter *s*. We focused our analysis on the information of the contrast and orientation of one image while keeping the other image at the same contrast and of an orthogonal orientation. The linear Fisher information were measured from the spike counts of the V4/MT excitatory neurons using a bias-corrected estimation (Kanitscheider et al. (2015); see Methods).

We find that neurons with stronger normalization encode more information per spike compared to neurons with weaker normalization (Figure 5). We grouped neurons based on their normalization indices and randomly sampled a various number of neurons within each group. We then divided the Fisher information of the sampled neurons by the their trial-averaged total number of spikes (See Methods). The Fisher information of contrast per spike is non-monotonic for neurons with weak normalization with a maximum at around 12 spikes and reduces to zero as the total number of spikes increases (Figure 5A1,A3). The variance of the Fisher information per spike across samples also peaks at around 10 spikes and largely shrinks when the total number of spikes is above 100. Neurons with stronger normalization encode more information per spike when the total number of spikes is below 100 (Figure 5A2). This is consistent with our observation in the previous section that the firing rates of neurons with strong normalization are sensitive to contrast changes in the images (Figure 4). The efficiency of information encoding of neurons with different normalization indices converges and decays to zero when the total number of spikes is large (Figure 5A3). The Fisher information of orientation shows similar trend as the information of contrast, except that the Fisher information per spike for orientation tends to decrease monotonically for neurons with weak normalization (Figure 5B).

**Figure 5.**
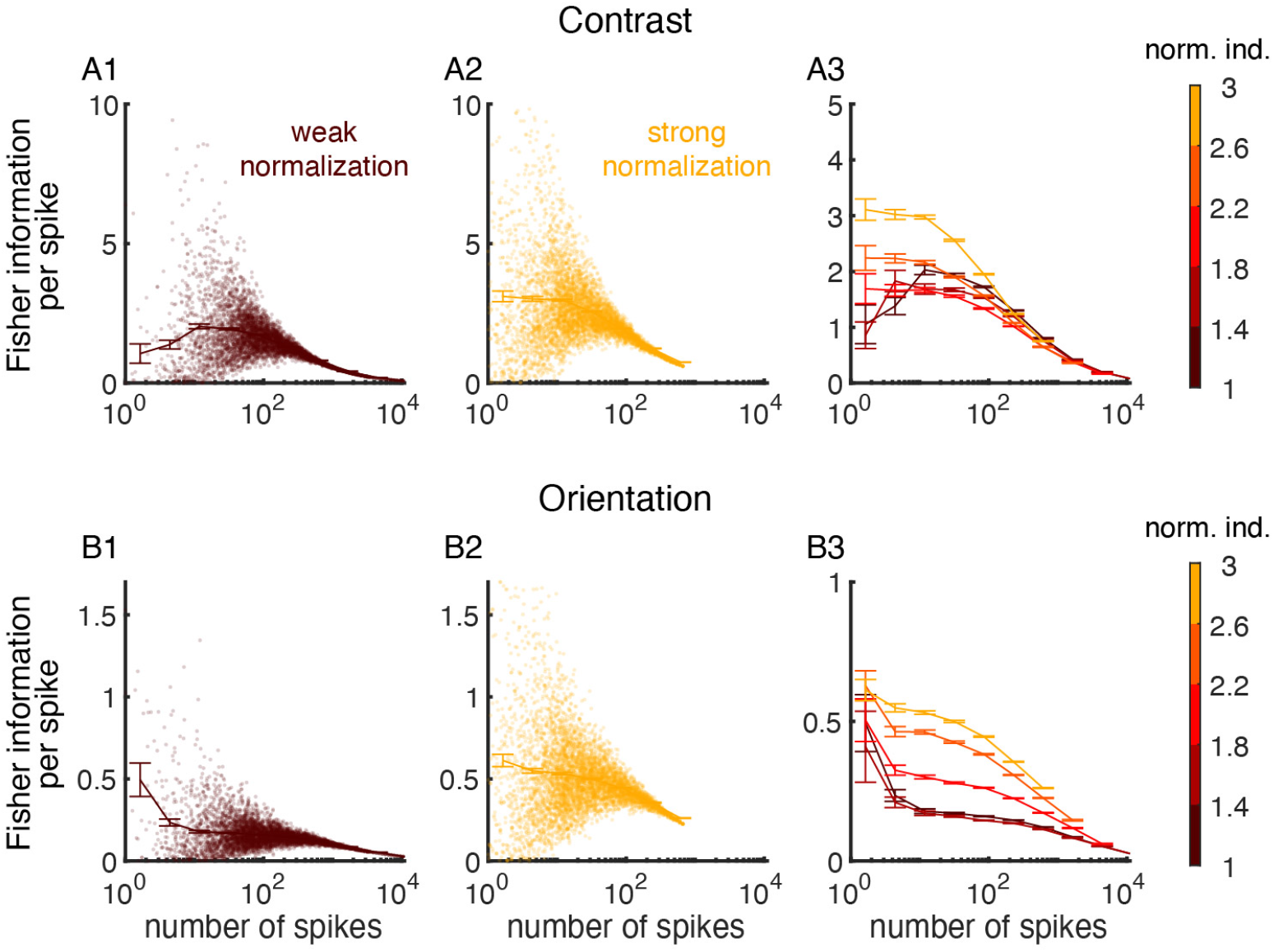
Neurons with stronger normalization encode information more efficiently. (**A**) The linear Fisher information (Eq. 2, see Methods) of the contrast of one of the two images per spike from neurons with different normalization indices. Various numbers of neurons were randomly sampled from the model V4/MT excitatory neurons whose normalization indices were within the specified range indicated by the color bar. The linear Fisher information of the sampled group of neurons was divided by the number of spikes from the sampled group of neurons during the time window used for calculating the Fisher information. (**A1**) The linear Fisher information per spike as a function of the total number of spikes from the sampled neurons. Neurons with normalization indices from 1 to 1.4 were sampled. Each dot is a different sampling of a group of neurons. The solid curves is the average for each bin of the number of spikes and the error bar is standard error. (**A2**) Same as **A1** except that neurons with normalization indices from 2.6 to 3 were sampled. (**A3**) The average Fisher information per spike as a function of the total number of spikes from the sampled neurons, for neurons of different ranges of normalization indices. Neurons with stronger normalization (lighter color) encode more information per spike than neurons with weak normalization (darker color). (**B1-3**) Same as **A1-3** for the linear Fisher information of the orientation of one of the two images.

### Heterogeneous normalization enhances the information of image contrast but not orientation

Lastly, we compare the information content in networks with different amount of heterogeneity in normalization. We have shown that the normalization index of a model neuron is strongly correlated with the inhibitory current it receives (Figure 3A-C). To reduce the heterogeneity in normalization, we constructed a control network with the same number of input connections (i.e. in-degree) to each V4/MT neuron, including both local and long-range connections, so that neurons receive roughly the same magnitude of currents. All the other parameters, such as the connection weights and the spatial spreads of connections were kept the same as the default network. Matching the in-degrees in a homogeneous network without spatial or tuning dependent connections leads to similar firing rates in neurons from the same cell type population (excitatory or inhibitory) (Brunel, 2000). Matching the in-degrees in our spatial network also largely reduces the spread of normalization indices (Figure 6A). The remaining heterogeneity in normalization is partly due to the tuning selectivity of neurons and the spatial arrangement of the pinwheel orientation map.

**Figure 6.**
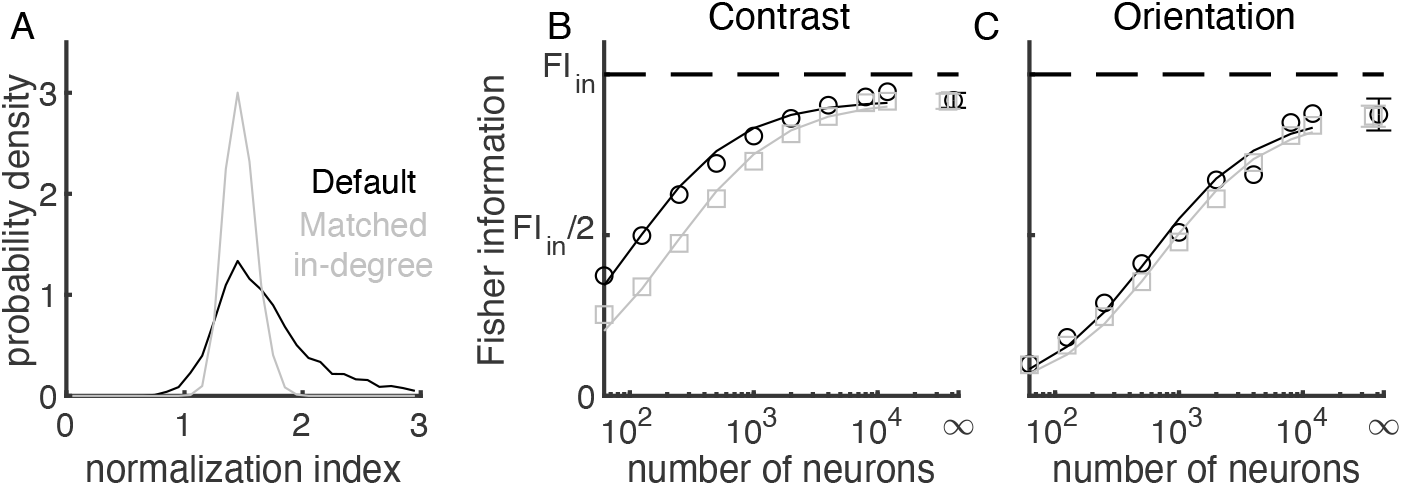
Heterogeneous normalization enhances the information of image contrast but not orientation. (**A**) The distribution of normalization indices is narrower in the network with matched in-degrees (grey) than that in the default network (black). In the network with matched in-degreess, the number of presynaptic neurons that project to each V4/MT neuron is fixed to be the same for each type of inputs (recurrent excitation, recurrent inhibition and feedforward excitation). The default network is the same as described in Figure 1 and analyzed in previous figures. (**B**) The linear Fisher information of the contrast of one image as a function of the number of excitatory neurons sampled from the model V4/MT network. Open circles are the numerical estimation of the linear Fisher information (Eq. 18). The asymptotic values of the linear Fisher information at limit of large number of sampled neurons (dots at *N* = ∞) are estimated by fitting Eq. 19, and solid curves are the fits (see Methods). Error bars are the 95% confidence intervals. Dashed line is the total amount of linear Fisher information from V1 neurons. The total input information is the same for both networks. (**C**) Same as **B** for the linear Fisher information of the orientation of one image.

As the number of sampled V4/MT neurons increases, the Fisher information of the sampled neurons increases and approaches the upper bound of the total input information from V1 neurons (Figure 6B,C). The information of image contrast is higher in the default network than that in the network with matched in-degrees, which has less heterogeneity in normalization (Figure 6B). The information difference between the two networks reduces as the number of sampled neurons increases and the information curves from both networks saturate to a similar level close to the upper bound of input information. This again demonstrates the efficiency of neural coding in networks with heterogeneity, that is, networks with more heterogeneity of normalization encode more information of contrast with a smaller number of neurons.

Interestingly, the information of orientation is very similar in both networks with different amounts of normalization heterogeneity across various number of sampled neurons, with the information from the default network being slightly higher (Figure 6C). This suggests that heterogeneous normalization is more beneficial for encoding contrast than orientation in our network.

## Discussion

Normalization mechanism has often been used to describe neurons’ sublinear responses to multiple stimuli in visual cortex (Britten and Heuer, 1999; Heeger, 1992; Ni et al., 2012; Rust et al., 2006; Verhoef and Maunsell, 2017)). However, neurons exhibit diverse response patterns in their integration of multiple stimuli; some neurons show facilitated responses to two stimuli, while some neurons are strongly suppressed by an addition of a non-preferred stimulus (Barbera et al., 2022; Guan et al., 2020; Ni et al., 2012; Ruff et al., 2016). The mechanism underlying the neuronal heterogeneity of normalization and its contribution to neural coding is not well studied. In this work, we analyzed response properties of a recurrent network of excitatory and inhibitory spiking neurons modeling the visual cortex. Our model neurons exhibit a range of normalization strengths that is consistent with experimental data (Ruff et al., 2016). We find that a neuron’s normalization strength is strongly correlated with the relative magnitude of inhibitory current it receives. In addition, our model reproduces the relationship between normalization strength and the spike count correlations between pairs of neurons observed in experimental data, which can be explained by the covariance with inhibitory current. Further, we demonstrate that model neurons with different normalization strengths and selectivity respond to different combinations of the stimulus contrasts. Model neurons with stronger normalization are more sensitive to the contrast difference of images and encode more stimulus information per spike. Lastly, we show that neuronal heterogeneity can be beneficial for coding, as networks with more heterogeneity encode more information of image contrast.

We find that strong recurrent coupling among the excitatory and inhibitory neurons and distinct sources of feedforward inputs are important for generating a wide range of heterogeneity of normalization strength. In networks with weak recurrent coupling, the distribution of normalization indices is narrow and the spike count correlations between neurons only weakly depend on their normalization strengths (Supp Fig S4). This suggests that strong recurrent connections amplify the heterogeneity in neuron responses to multiple inputs. In networks with strong recurrent coupling and disordered connections (i.e. no spatial or tuning dependence of connections), neurons exhibit a range of normalization strengths when the network receives two sources of feedforward inputs (Supp Fig S3). The distribution of normalization strengths broadens as the two sources of inputs target more distinct populations of neurons. The range of normalization strengths is similar to that of the detailed circuit model of visual cortex where the two images of orthogonal orientations activate largely non-overlapped groups of neurons in the V4/MT network (Figure 1). Our results are consistent with a recent finding where models with strong recurrent coupling explains the large distribution of rate changes in monkey visual cortex induced by optogenetic stimulation (Sanzeni et al., 2023).

The balanced network model (Van Vreeswijk and Sompolinsky, 1996) has been successful at explaining the genesis of neuronal variability and the close relationship of excitation and inhibition observed in experiments in cortex (Haider et al., 2006; Okun and Lampl, 2008; Xue et al., 2014). However, balanced networks produce a linear relationship between input and output rate, which has been considered as a limitation for complex computation (van Vreeswijk and Sompolinsky, 1998). Recent theoretical work suggests that nonlinear computation can be achieved in balanced networks when some neurons are silenced by excess inhibition, creating a local imbalance in currents (Baker et al., 2020). They show that a network can produce sub-linear summation, if individual stimulus silences a group of neurons when presented alone (Baker et al., 2020). Our finding is different from this work in that we analyzed neurons with positive rates in all three stimulus conditions (stimulus 1 alone, stimulus 2 alone or both). In other words, neurons in our model do not need to be silenced in one stimulus condition to show sublinear summation. Consistent with the previous work, our results demonstrate that individual neurons can exhibit diverse and nonlinear response functions to multiple stimuli, even though the population averaged rate remains mostly linear (Figure 4A1-4).

Even though the normalization phenomena has been widely observed in visual cortex, as well as in other brain regions, the source of normalization has been under debate. It was initially hypothesized that the divisive form of normalization can be implemented by shunting inhibition (Carandini et al., 1997). However, it has been shown that shunting inhibition alone results in a subtractive, not divisive, modulation of firing rates (Chance et al., 2002). Instead, divisive gain modulation can be achieved by modulating both excitatory and inhibitory inputs in a balanced manner (Chance et al., 2002). Later experimental findings suggest that it is the excitation, rather than inhibition, that underlies the sublinear responses of neurons since both excitation and inhibition are suppressed with an addition of non-preferred stimulus (Sato et al., 2016). Recent experiments suggest that a feedforward mechanism is sufficient to account for the sublinear responses of neurons in primary visual cortex without invoking a recurrent mechanism (Barbera et al., 2022; Priebe and Ferster, 2006), though mechanisms for higher-order visual areas are not studied. In our model, we find that the normalization strength of a neuron is highly anti-correlated with the inhibitory current it receives, and is only weakly correlated with the feedforward and recurrent excitatory currents (Figure 3A-C). Moreover, the covariance with the inhibitory current also explains the relationship between spike count correlation and normalization. Neurons with stronger normalization receive relatively more inhibitory currents, which cancel out more correlation in their currents, making those neurons less correlated with other neurons. Our results are consistent with previous theoretical work which suggests that the population firing rate patterns in cortical circuits are primarily determined by inhibitory currents (Mongillo et al., 2018). Together, our results emphasize the role of inhibition in determining the normalization strength and neuronal correlations.

Numerous experimental work has demonstrated that the neural mechanisms of normalization and selective attention are closely related (Carandini and Heeger, 2012; Ni et al., 2012; Reynolds and Heeger, 2009; Reynolds et al., 1999; Treue and Maunsell, 1996). In particular, the neuronal heterogeneity of attentional modulation in firing rates is highly correlated with the neuronal heterogeneity of normalization, meaning that neurons that demonstrate stronger normalization are also more modulated by attention (Lee and Maunsell, 2009; Ni et al., 2012). In addition, the spike count correlations among neurons with stronger normalization are also more modulated by attention (Verhoef and Maunsell, 2017). Both experimental and modeling work suggests that inhibitory neurons are more targeted by attention than excitatory neurons, which could stabilize population dynamics and reduce neural correlations (Huang et al., 2019; Kanashiro et al., 2017; Mitchell et al., 2007; Thiele et al., 2016). Our finding of the strong correlation between normalization strength and inhibitory current (Figure 3C) suggests that inhibition may be the unifying mechanism that relates the neuronal heterogeneity of normalization and attentional modulation. Future extension of our model is needed to explore the interplay between normalization and attention mechanisms in enhancing the neural representation of attended stimuli.

Several network models have been proposed to explain the normalization mechanism (Heeger and Zemlianova, 2020; Rubin et al., 2015; Somers et al., 1998). One of them, the stabilized supralinear network (SSN) model (Hennequin et al., 2018; Rubin et al., 2015), is closely related to our model. The SSN model also has strong recurrent excitation which is stabilized by inhibitory feedback and the recurrent connections depend on tuning similarity. The SSN model reproduces the sublinear summation of neuronal responses to two stimuli and the quenching of neuronal variability by stimulus contrast. The key differences between our model and the SSN model are that our model consists of spiking neurons instead of rate units and that the population rate of our model do not show strong saturation as input strength increases. Nevertheless, both our model and the SSN model may share common mechanisms for generating heterogeneous normalization strength. For comparison, we implemented the two-dimensional version of the SSN model with probabilistic connections to introduce heterogeneity (Supp Fig S8; Rubin et al. (2015)). We found that the normalization index of a neuron strongly depended on the neuron’s preferred orientation in the SSN model, which was not observed in our experimental data and the spiking network model (compare Supp Fig S8B with Supp Fig S2). There was a similar relationship between spike count correlations and normalization indices in the SSN model, however, the relationship was absent after we matched for the distribution of tuning preferences across normalization indices (Supp Fig S8C).

Past work has proposed several computational benefits of normalization (summarized in review Carandini and Heeger (2012)). For example, by adapting neurons’ response range based on background input, neurons can remain sensitive to small changes in stimulus. This is most evident for retinal neurons which need to respond to light intensities over a range of several orders of magnitude (Rieke and Rudd, 2009). In visual cortex, it has been shown that divisive normalization can reduce the statistical redundancy present in natural images (Schwartz and Simoncelli, 2001). Divisive normalization can also implement marginalization in the framework of probabilistic population code (Beck et al., 2011). Complementary to these works, our results reveal additional benefits of normalization, that is, neurons with stronger normalization encode more stimulus information per spike (Figure 5).

In addition, we demonstrate that the neuronal heterogeneity of normalization contributes to coding. Past works have shown that cellular heterogeneity, such as in spiking threshold or excitability, can increase network responsiveness (Di Volo and Destexhe, 2021), improve network resilience to changes in modulatory inputs (Hutt et al., 2023) and enhance the mutual information between stimulus and neural responses (Kastner et al., 2015; Sharpee, 2017). Heterogeneity in neuronal time scales can also improve learning in tasks with rich intrinsic temporal structure (Perez-Nieves et al., 2021). Our work is different from these works in that our model neurons are homogeneous in terms of their cellular properties, and the heterogeneity in their response properties is generated by network interactions. The heterogeneous normalization strength allows the neural population to encode different contrast combinations of the two images (Figure 4). In networks with less heterogeneity of normalization, more neurons are needed to encode the same amount of information (Figure 6B). Therefore, neuronal heterogeneity in normalization improves the efficiency of information coding. Interestingly, we find that heterogeneity has a larger impact on the information of image contrast than on the information of orientation. Future work is needed to investigate how neuronal heterogeneity impacts population representational geometry of multiple stimulus features in circuit models.

## Methods

### Spiking neuron network model of visual cortex

The model consists of two layers of spiking neurons, modeling the primary visual cortex (V1) and a higherorder visual area (V4 or MT), respectively (Figure 1A). Neurons from the two layers are arranged on a uniform grid covering a a rectangle of size Γ = [0, 2] × [0, 1]. There are 5,000 excitatory neurons in the V1 layer, 20,000 excitatory neurons and 5,000 inhibitory neurons in the V4/MT layer.

#### V1 model neurons

V1 neurons are modeled as linear-nonlinear Poisson spiking neurons, following previous models (Huang et al., 2022; Kanitscheider et al., 2015). V1 neurons are divided into two populations, V1_1_ and V1_2_, each of which has a non-overlapping receptive field centering on each Gabor image, respectively. Neurons located at the left half of the rectangle Γ ([0, 1] × [0, 1]) have receptive fields centered at and with the same size as image 1, while those located at the right half of the rectangle ([1, 2] × [0, 1]) have receptive fields centered at and with the same size as image 2. The receptive field of a neuron from population *k* (*k* = 1, 2) is modeled as a Gabor filter:

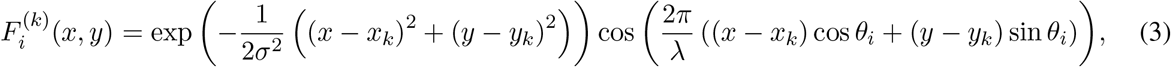

where the subscript *i* denotes the neuron’s index, *σ* = 0.2 is the standard deviation of the Gaussian envelope, *λ* = 0.6 represents the wavelength of the sinusoidal factor, *x* and *y* are the coordinates of the neuron, (*x*_*k*_, *y*_*k*_) is the center of the receptive field ((*x*_1_, *y*_1_) = (0.5, 0.5) for V1_1_ neurons and (*x*_2_, *y*_2_) = (1.5, 0.5) for V1_2_ neurons) and *θ*_*i*_ is the preferred orientation of neuron *i*. The preferred orientation of each neuron was assigned according to a pinwheel orientation map generated with the method from Kaschube et al. (2010) (Supp Materials Eq. 20). The preferred orientation at (*x, y*) is *θ*_*i*_(*x, y*) = angle(*z*(*x, y*))*/*(2*π*) and

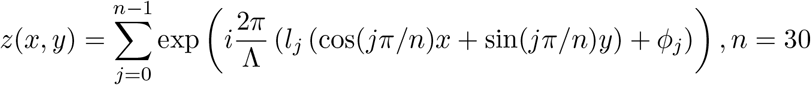

where Λ = 0.125 is the average column spacing, *l*_*j*_ = ±1 is a random binary vector and the phase *ϕ*_*j*_ is uniformly distributed in [0, 2*π*].

Spike trains of V1 neurons are generated as inhomogeneous Poisson process with instantaneous rate

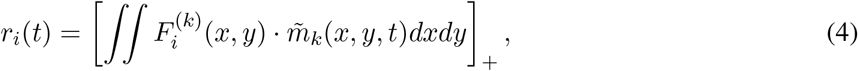

where 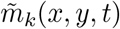 is the pixel value of image *k* (*k* = 1, 2) defined below (Eq. 6) and 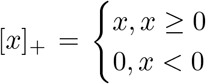 denotes half rectification. In the presence of image *k*, the average of rate of V1_*k*_ neurons was 10 Hz. In the absence of image *k*, V1_*k*_ neurons had a spontaneous rate of 5 Hz.

Two Gabor images of orthogonal orientations are presented to the V1 neurons, either individually or simultaneously. Each image has 25 × 25 pixels and is defined as

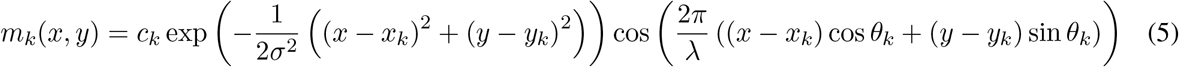

where *c*_*k*_ is the contrast of image *k*, and *σ* and λ are the same as the Gabor filters of the V1 neurons (Eq. 3). Each pixel is corrupted by independent additive noise as

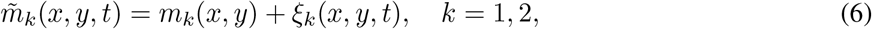

where *ξ*_*k*_(*x, y, t*) is modeled as Ornstein-Uhlenbeck process

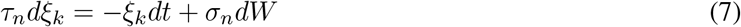

with *τ*_*n*_ = 40 ms, *σ*_*n*_ = 3.5 and *dW* being a Wiener process.

#### V4/MT model neurons

V4/MT layer is a recurrently coupled network of excitatory (*α* = e) and inhibitory neurons. The neuronal and synaptic parameters are the same as in our previous model (Huang et al., 2019). The preferred orientation of each neuron was assigned with a separately generated orientation map using the same method as that of V1 neurons. Each neuron is modeled as an exponential integrate-and-fire (EIF) neuron whose membrane potential is described by:

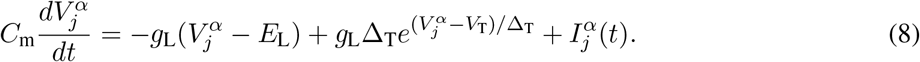

A spike is generated each time 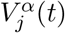 exceeds a threshold, *V*_th_. Then the neuron’s membrane potential is held for a refractory period, *τ*_ref_, after which it is reset to a fixed value *V*_re_. Neuron parameters for excitatory neurons are *τ*_m_ = *C*_m_*/g*_L_ = 15ms, *E*_L_ = − 60mV, *V*_T_ = − 50mV, *V*_th_ = − 10mV, Δ_T_ = 2mV, *V*_re_ = − 65mV and *τ*_ref_ = 1.5ms. Inhibitory neurons are the same except *τ*_m_ = 10ms, Δ_T_ = 0.5mV and *τ*_ref_ = 0.5ms. The total current to the *j*^th^ neuron is:

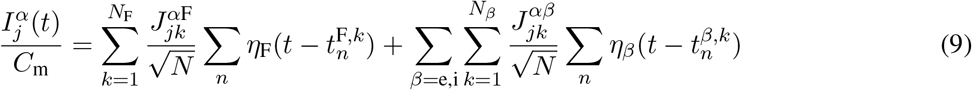

where *N* = *N*_e_ + *N*_i_ is the total number of the network population. The postsynaptic current is given by

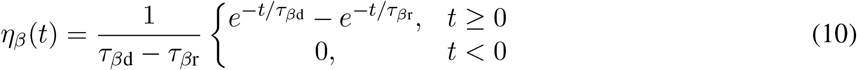

where *τ*_er_ = 1ms, *τ*_ed_ = 5ms for excitatory synapses and *τ*_ir_ = 1ms, *τ*_id_ = 8ms for inhibitory synapses. The feedforward synapses from V1 neurons to V4/MT neurons have a fast and a slow component.

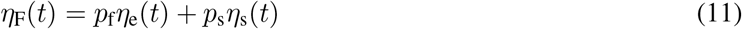

with *p*_f_ = 0.2, *p*_s_ = 0.8. *η*_s_(*t*) has the same form as equation 10 with a rise time constant 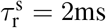 and a decay time constant 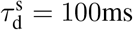.

#### Network connections

There are two types of feedforward and recurrent excitatory connections projecting to the excitatory V4/MT neurons. 85% of connections are generated according to connection probability that depends only on the physical distance between neurons (Eq. 12). The remaining 15% of excitatory connections are randomly chosen from similarly tuned neurons and do not depend on space. The probability of inhibitory projections and the projections to the inhibitory neurons only depends on distance (Eq. 12) and not on tuning similarity.

The distance-dependent connections are sampled according to probability function, *p*_*αβ*_(**x**_1_, **x**_2_), between a neuron from population *β* at location **x**_1_ = (*x*_1_, *y*_1_) to a neuron from population *α* at location **x**_2_ = (*x*_2_, *y*_2_), *α, β* ={e, i}.

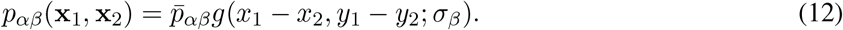

Here 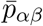 is the mean connection probability and

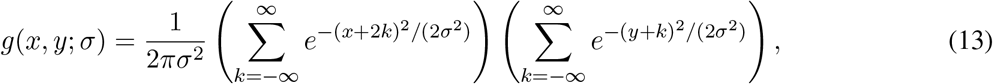

is a wrapped Gaussian distribution periodic on the domain Γ. The excitatory and inhibitory recurrent connection widths of the V4/MT layer were *σ*_e_ = *σ*_i_ = 0.2 and the feedforward connection width from the V1 layer to the V4/MT layer was *σ*_ffwd_ = 0.1. A presynaptic neuron was allowed to make more than one synaptic connection to a single postsynaptic neuron.

The long-range excitatory connections are sampled between similarly tuned neurons, i.e. cos (*θ*_*i*_ − *θ*_*j*_) ≥ 0.6, where *θ*_*i*_ and *θ*_*j*_ are the preferred orientations of neuron *i* and *j*.

The recurrent synaptic weights within the V4/MT layer were *J*_ee_ = 80m<u>V</u>, *J*_ei_ = − 240mV, *J*_ie_ = 40Mv and *J*_ii_ = − 300mV. Note that each synaptic weight was scaled by 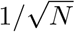 (Eq. 9). The mean connection probabilities were 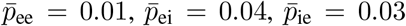 and 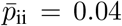.The out-degrees were 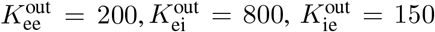, and 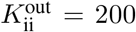.The feedforward connection strengths from V1 layer to V4/MT layer were *J*_eF_ = 160mV and *J*_iF_ = 140mV, with probabilities 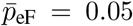 and 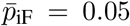 (out-degrees 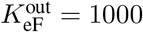 and 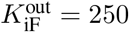).

#### Network with matched in-degrees

In the network with matched-in-degrees (Figure 6), the number of presynaptic neurons (in-degrees) that project to each V4/MT neuron was matched to be the same across each population for each type of connections (V4/MT excitatory, V4/MT inhibitory, or V1 excitatory), while the out-degrees were allowed to vary. All other parameters including synaptic weights and connection probabilities were the same as the default network.

#### Simulation

All simulations were performed on the CNBC Cluster in the University of Pittsburgh. All simulations were written in a combination of C and Matlab (Matlab R 2021b, Mathworks). The differential equations of the neuron model were solved using forward Euler method with time step 0.05 ms.

### Datasets and analysis

Neuronal activity was collected from four adult male rhesus monkeys (*Macaca mulatta; monkeys BR, JD, ST, SY*) as they were passively fixating at superimposed orthogonal drifting gratings at a range of contrasts (details are described in Ruff et al. (2016)). All animal procedures were approved by the Institutional Animal Care and Use Committees of the University of Pittsburgh and Carnegie Mellon University.

MT data was collected with 24-channel V-Probes and 24-channel linear microarrays in area MT of two monkeys (Figure 2B). There were a total of 2,133 visual stimulus trials for 769 units and 10,600 pairs from 28 recording sessions. V4 data was collected with a pair of 6 × 8 microelectrode arrays implanted in area V4 of two monkeys (Figure 2C). In one monkey, both arrays were in V4 in the right hemisphere. In the other monkey, arrays were implanted bilaterally in area V4. There were a total of 2,160 visual stimulus trials for 1,276 units and 39,719 pairs from 21 recording sessions. V1 data was collected with a 10 × 10 microelectrode array implanted in area V1 of two monkeys (Figure S5). There were a total of 1,467 visual stimulus trials for 2,124 units and 97,169 pairs from 23 recording sessions.

In our analysis, we included units if their response to 0% contrast stimuli was significantly different from the average response to stimuli with at least 50% contrast (*t*-test, *P* < 0.01). Pairs of units that came from the same electrode were excluded for correlation analysis. For each unit recorded in each stimulus condition, spike counts were calculated from 30 to 230 ms for V1 units and from 50 to 250 ms for V4 and MT units after stimulus onset, to allow for latency in response. Each stimulus was presented for 200 ms. We quantified spike count correlations as the Pearson’s correlation coefficient between spike count responses to repeated presentations of the same stimulus. This measure is extremely sensitive to outliers, so we did not analyze trials for which the response of either unit was more than three standard deviations away from its mean (following the convention of Kohn and Smith (2005)).

To compute the normalization index of recorded units, we included the mean spike counts in response to stimulus conditions where the contrast of one stimulus was 50% and the other was 0%, and where the contrast of both stimuli were 50%. The normalization index was computed using Equation 14 for each combination of orthogonal drifting gratings and then averaged across all combinations within a session. To compute the selectivity of recorded units, we included the mean spike counts in response to stimulus conditions where the contrast of one stimulus was 50% and the other was 0%. Tuning similarity was quantified as the Pearson’s correlation coefficient between mean spike count responses to each stimulus direction presented alone, with 50% contrast and the contrast of the other direction equal to 0%. The spike count correlation of a pair of units in one session was averaged across the stimulus conditions used to compute the normalization index. In Figure 2B, 2C, and S5, the spike count correlation was averaged across unit pairs from all recording sessions.

### Statistical methods

#### Normalization index

The normalization index of a neuron is defined as the sum of a neuron’s firing rate to each one of the two stimuli when presented alone divided by its firing rate when both stimuli are presented togenther (Ruff et al., 2016):

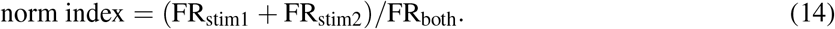

A normalization larger than 1 indicates sublinear summation.

The current normalization index is defined similarly:

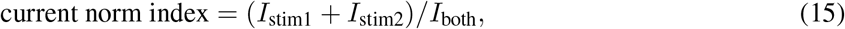

where *I* is the average recurrent excitatory, recurrent inhibitory or feedforward excitatory current a neuron receives.

#### Spike count correlation and current covariance

We computed the spike count correlation of V4/MT model neurons when both images of orthogonal orientations were presented together. Spike counts were computed using a sliding window of 200 ms with 50 ms step size and the Pearson correlation coefficients were computed between pairs of neurons. In Figure 2A, there were 10 simulations and each simulation was 20 seconds long. The first 1 second of each simulation was excluded from the correlation analysis to avoid transient effects. Neurons whose average firing rates were within one standard deviation from the mean population rates in all of the three stimulus conditions (image 1 alone, image 2 alone, or both images presented together) were included. In total, 8663 excitatory V4/MT neurons were sampled to compute spike count correlations.

To compute current covariance (Figure 3D,E), each type of synaptic currents (feedforward excitation, recurrent excitation and recurrent inhibition) to each neuron were recorded every 10 ms. The excitatory current combines both feedforward and recurrent excitation. In total, 4163 neurons were sampled to compute current covariance. The simulation was 20 seconds long.

#### Tuning similarity

Tuning similarity between a pair of neurons is defined as the Pearson correlation between their tuning curves of orientation:

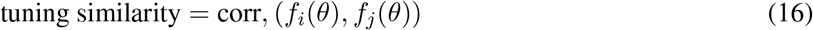

where *f*_*i*_ and *f*_*j*_ are neuron *i* and neuron *j*’s tuning curves, respectively. Tuning curves were computed using one image.

#### Selectivity

Selectivity measures how selective a neuron is to stimulus 1 compared to stimulus 2.

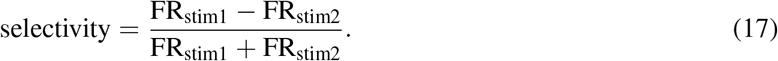

Selectivity takes a value between −1 and 1. The larger the absolute value of selectivity is, the more selective the neuron is to its preferred stimulus. A selectivity equaling 0 means there is no preference of the neuron between the two stimuli.

#### Linear Fisher information

To compute the linear Fisher information, stimulus 2 was presented during 200 ms intervals (ON) interleaved with 300 ms OFF intervals, during which the spike trains of V1_2_ neurons were independent Poisson process with rate 5 Hz. Meanwhile stimulus 1 was present throughout a simulation. Spike counts of V4/MT excitatory neurons during the ON intervals were used to compute the linear Fisher information. Each simulation was 20 seconds long. The first spike count in each simulation was excluded. The connectivity matrices were fixed for all simulations and the initial state of each neuron’s membrane potential was randomized in each simulation.

For the linear Fisher information of contrast, the contrast of stimulus 2 during ON intervals was randomly chosen from *c*_1_ = *c* + *δc/*2 and *c*_2_ = *c* − *δc/*2, where *c* = 0.5 and *δc* = 0.01. For the linear Fisher information of orientation, the orientation of stimulus 2 during ON intervals was randomly chosen from *θ*_1_ = *θ* + *δθ/*2 and *θ*_2_ = *θ* − *δθ/*2, where *θ* = 0.5*π* rad and *δθ* = 0.02 rad. The linear Fisher information of V4/MT neurons was computed using a bias-corrected estimate (Kanitscheider et al. (2015))

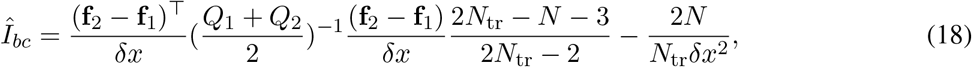

where *δx* = *δc* or *δθ*, **f**_*i*_ and *Q*_*i*_ are the empirical mean and covariance, respectively, for *c*_*i*_ or *θ*_*i*_. *N*_tr_ is the number of trials for each *c*_*i*_ or *θ*_*i*_.

We used the fitting algorithm proposed by Kafashan et al. (2021) to estimate the asymptotic value of the linear Fisher information, *I*_∞_, at the limit of *N*→ ∞, where *N* is the number of sampled neurons (Figure 6B,C). Briefly, the theory of information-limiting correlations (Moreno-Bote et al. (2014)) shows that the linear Fisher information, *I*_*N*_, in a population of *N* neurons can be decomposed into a limiting component, *I*_∞_, and a nonlimiting component *I*_0_(*N*),

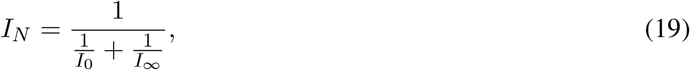

where we assume that the nonlimiting component increases linearly with *N*, i.e., *I*_0_ = *aN*. Hence, Equation 19 can be rewritten as

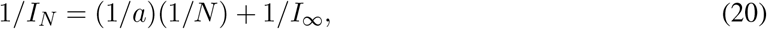

which shows that 1*/I*_*N*_ scales linearly with 1*/N* with 1*/I*_∞_ as the intercept. Hence, we do a linear fit of 1*/I*_*N*_ versus 1*/N*, with *N* varying from 8 to 12000 and estimate 1*/I*_∞_.

The linear Fisher information from V1 neurons can be estimated analytically as ((Kanitscheider et al., 2015),

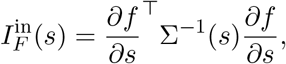

where *s* = *c* or *θ*, with

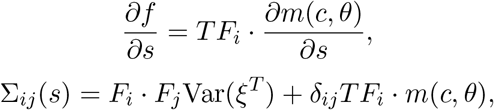

where 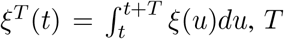, is the time window for spike counts, and *δ*_*ij*_ is a Kronecker delta, which is 1 if *i* = *j*, and 0 otherwise. We can calculate the variance of the integrated noise over time window *T* as 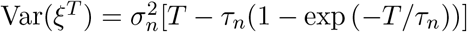.

In Figure 5, a number of neurons were randomly sampled from the model V4/MT excitatory neurons whose normalization indices were within a specified range. The number of sampled neurons, *N*, was logarithmically spaced between 1 and 8103. For *N* < 545 neurons, the number of samplings for each *N* decreased from 200 to 14 proportionally with *log*(*N*), rounded to the nearest integer. For *N* ≥545 neurons, there were 5 samplings for each *N*. The linear Fisher information of the sampled group of neurons was divided by the average number of spikes from the sampled group of neurons during the time window used for calculating the Fisher information (*T* = 200 ms). In Figure 6B,C, the number of neurons were *N* = 8, 16, 31, 62, 125, 250, 500, 1000, 2000, 4000, 8000, 12000. There were 20 samples of neurons for each *N*. Neurons with firing rates less than 1 Hz were excluded. There were 304,000 spike counts in total for *c*_1_ and *c*_2_, or *θ*_1_ and *θ*_2_ conditions.

## Acknowledgments

We were supported by National Institutes of Health grants RF1NS121913 (C.H., M.C.), R01EY022930 (M.C.) and R01EY034723 (M.C.), Simons Collaboration on the Global Brain awards NC-GB-CULM-00002794-06 (C.H) and 542961SPI (M.C.), NSF CAREER award 2337640 (C.H.) and the University of Pittsburgh (C.H.).

## Author Contributions

D.S. and C.H. conceived the project; D.S. performed the simulations and analysis, in consultation with C.H. and M.C.; D.R. and M.C. provided the experimental data; C.H. supervised the project; all authors contributed to writing the manuscript.

## Data and Software Availability

Computer code for all simulations and data analysis will be available online upon publication.

## Supplementary Information

**Figure S1:**
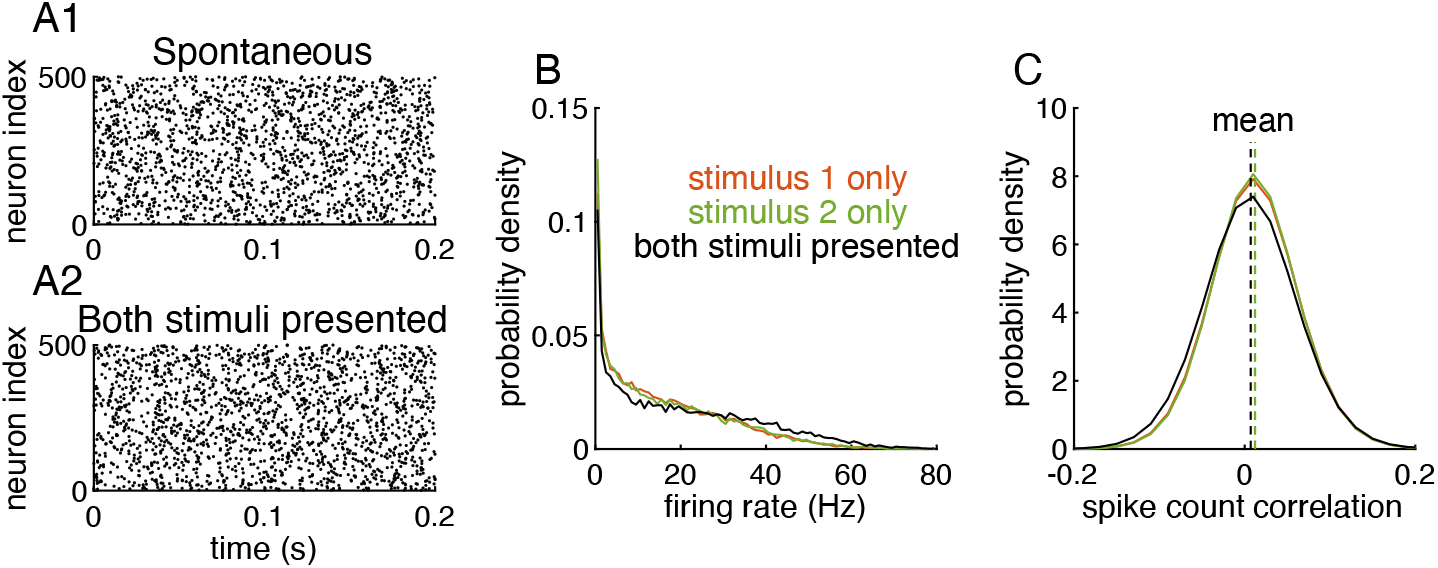
Related to Figure 1. The V4/MT network activity is stable and asynchronous in both spontaneous and evoked states. (**A1**) Spike raster of V4/MT neurons when both V1_1_ and V1_2_ neurons have homogeneous rates of 10 Hz. 500 neurons were randomly sampled from the V4/MT population. (**A2**) Spike raster of V4/MT neurons when two images are simultaneously presented. (**B**) The firing rate distributions of V4/MT neurons when only stimulus 1 (red), only stimulus 2 (green) or both stimuli are presented (black). (**C**) The distributions of spike count correlations of V4/MT neuron pairs from the three stimulus conditions. Dashed lines indicate the average spike count correlations (0.012 when only stimulus 1 is presented, 0.012 when only stimulus 2 is presented, 0.007 when both stimuli are presented).

**Figure S2:**
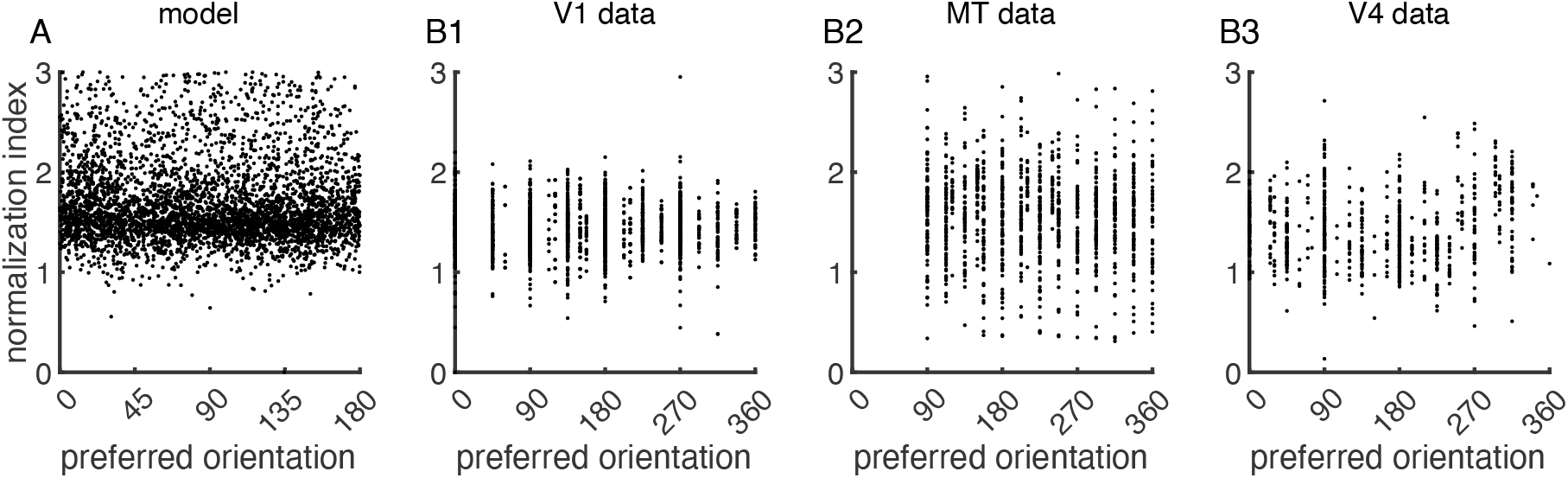
Related to Figure 2. The normalization index of a neuron is independent from its tuning preference in both the model and data. (**A**) The normalization index and the tuning preference of model V4/MT neurons are statistically independent. We use permutation test to assess the statistical significance of the null hypothesis that the normalization index and tuning preference are independent. First, we calculate the mutual information between the normalization index and tuning preference of V4/MT neurons. Then, we shuffle the normalization index of neurons 2000 times and recalculate the mutual information for each permutation. Lastly, we compute the p-value as the proportion of permutations with higher mutual information than that of the original observation. There is no significant dependence between the normalization index and the tuning preference of model V4/MT neurons (*p* = 0.31). (**B**) The normalization index and the tuning preference of experimentally recorded neurons are statistically independent in V1 area (**B1**, *p* > 0.05 for all 23 recording sessions), MT area (**B2**, *p >* 0.05 for 21 out of 28 recording sessions), and V4 area (**B3**, *p* > 0.05 for 14 out of 21 recording sessions).

**Figure S3:**
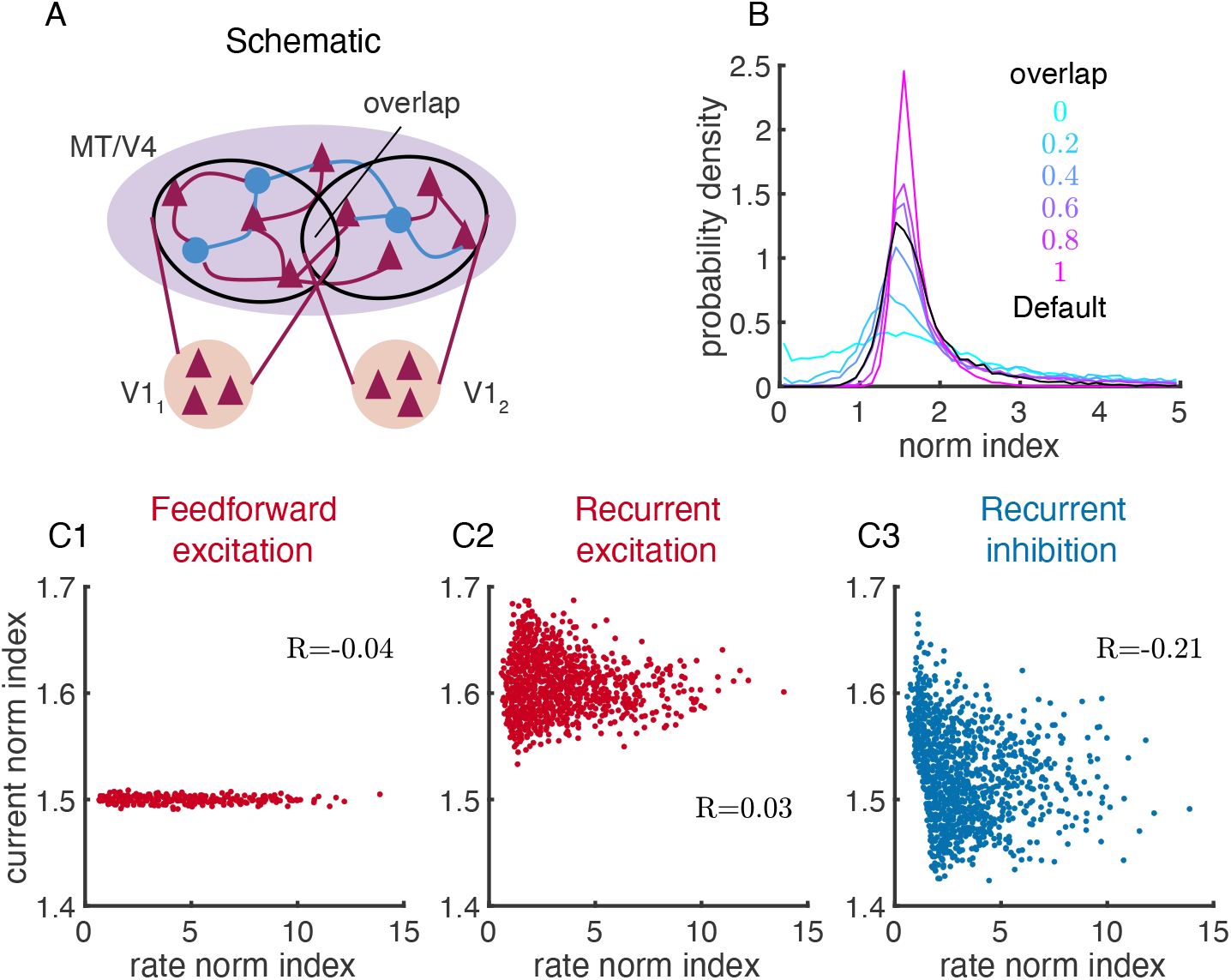
Related to Figures 1 and 3. Neurons exhibit a range of normalization strength in networks with strong recurrent coupling and disordered connections. (**A**) The schematic of the disordered network. The excitatory and inhibitory connections within V4/MT layer are random and with the same connection probability as those of the default model with spatial structure. The two V1 populations project randomly to the excitatory and inhibitory neurons in the V4/MT layer with the same connection probabilities as those in the default model. A fraction, *α*, of neurons in the V4/MT layer can receive input from both V1 populations, which represents the overlap of the feedforward projections from V1_1_ and V1_2_. The spike trains of V1 neurons are independent Poisson processes with fixed rates. In Stimulus 1 alone condition, V1_1_ neurons have firing rate of 10Hz and V1_2_ neurons have rate 5Hz. In Stimulus 2 alone condition, V1_1_ neurons have firing rate of 5Hz and V1_2_ neurons have rate of 10Hz. When both stimuli are presented, both V1_1_ and V1_2_ neurons fire at 10 Hz. (**B**) The distribution of normalization indexes is broader in networks with smaller overlap, *α*. The distribution of normalization indexes from the default network (black; same as that in Figure 1C) is similar to that of the random network model with *α* = 0.6. The normalization index of neurons in the random network model is calculated in the same way as in the default model, i.e. norm index = (FR_stim1_ + FR_stim2_)*/*FR_both_. (**C**) Relationship between current and rate normalization indexes in a random network model with *α* = 0.6. Only the normalization index of recurrent inhibitory current has a strong correlation with that of the firing rate. The range of the normalization index of feedforward excitatory current is very small in the random network model, since there is no spatial or tuning dependent connections.

**Figure S4:**
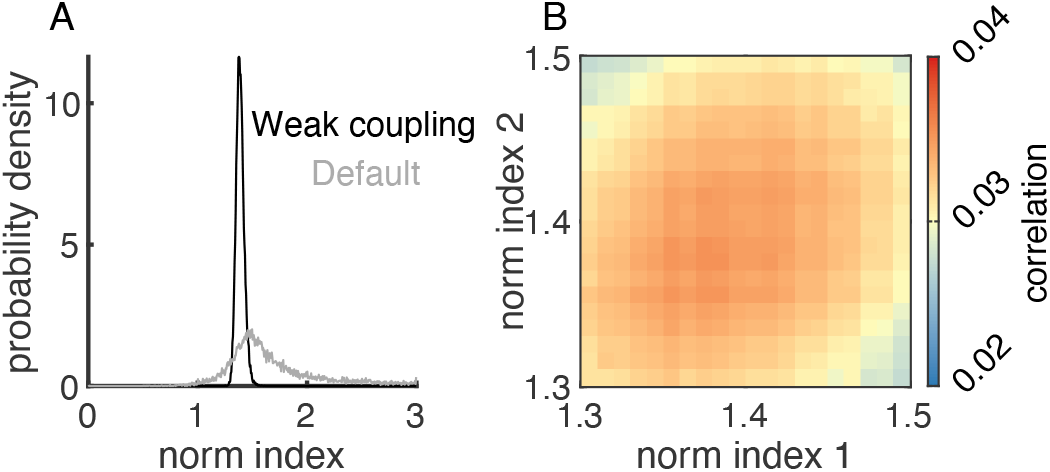
Related to Figures 1 and 2. Networks with weak recurrent coupling produce homogeneous normalization strength and weak dependence of spike count correlations on normalization. (**A**) The distribution of normalization indexes is much narrower in a network with weak recurrent coupling (black) compared to that in the default network (gray). The gray curve is the same as in Figure 1C but with different scale of the y-axis. (**B**) The dependence of spike count correlations on normalization index was weaker in the network with weak recurrent coupling. The range of spike count correlations between pairs of model V4/MT neurons was 0.008 in the network with weak recurrent coupling, compared to a range of 0.02 in the default network (Figure 2A1). In the network with weak recurrent coupling, the strengths of all recurrent connections were decreased to 10% of those in the default network. Independent white noise currents with mean 0 and standard deviation 6.8 mV/ms were applied to every excitatory and inhibitory V4/MT neuron such that the neurons’ f-I curve was the same as that in the default network. The strengths of feedforward projections were scaled down by a factor of 2.9 to keep the mean firing rate of V4/MT neurons the same as that in the default network. The default network was the same as that used in Figures 1 and 2.

**Figure S5:**
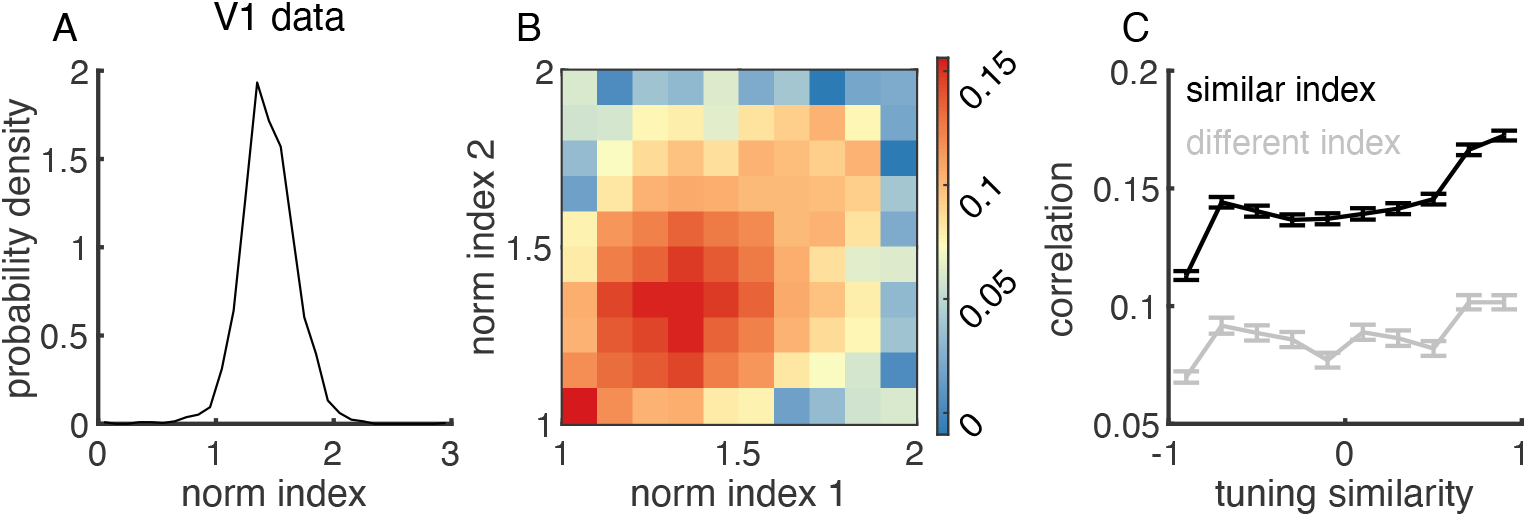
Related to Figure 1 and 2. Heterogeneity of normalization index and normalization-related modulation of spike count correlations are also observed in experimental data recorded from V1 area. (**A**) The recorded V1 neurons exhibit a similar range of normalization indexes as V4 and MT neurons (Figure 1D). (**B**) Spike count correlations between recorded V1 neurons depend on their normalization indexes. Same format as Figure 2A1,B1,C1). (**C**) Across all levels of tuning similarity, the spike count correlations between recorded V1 neurons with similar normalization indexes are consistently larger than those of neurons with distinct normalization indexes. Same format as Figure 2A4,B4,C4). Data from Ruff et al., 2016.

**Figure S6:**
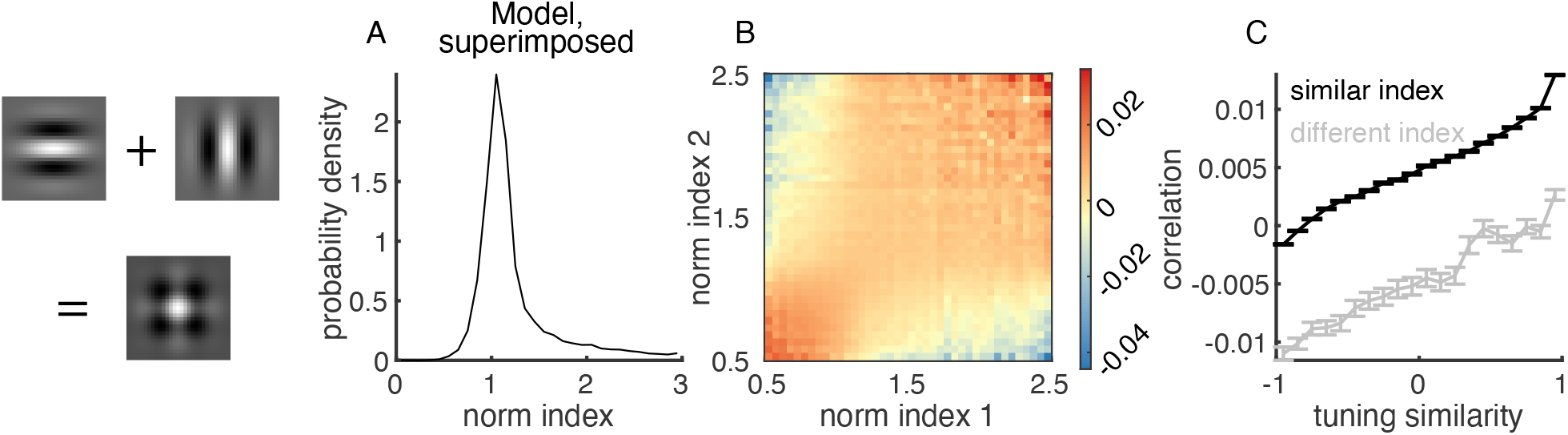
Related to Figures 1 and 2. Heterogeneity in normalization index and normalization-related modulation of spike count correlations in the model in response to two superimposed Gabor images. (**A**) Distribution of normalization indexes (Eq. 14 where FR_both_ was the firing rate when two superimposed Gabor images were presented. (**B**) Spike count correlations as a function of the normalization indexes of the two neurons in the pair. (**C**) Neurons with similar normalization indexes have higher noise correlations than those with different normalization indexes. This relationship is consistent across neuron pairs with various tuning similarities. Model results with superimposed Gabor images are qualitatively the same as those with two separate Gabor images (Figures 1C, 2A1,A4).

**Figure S7:**
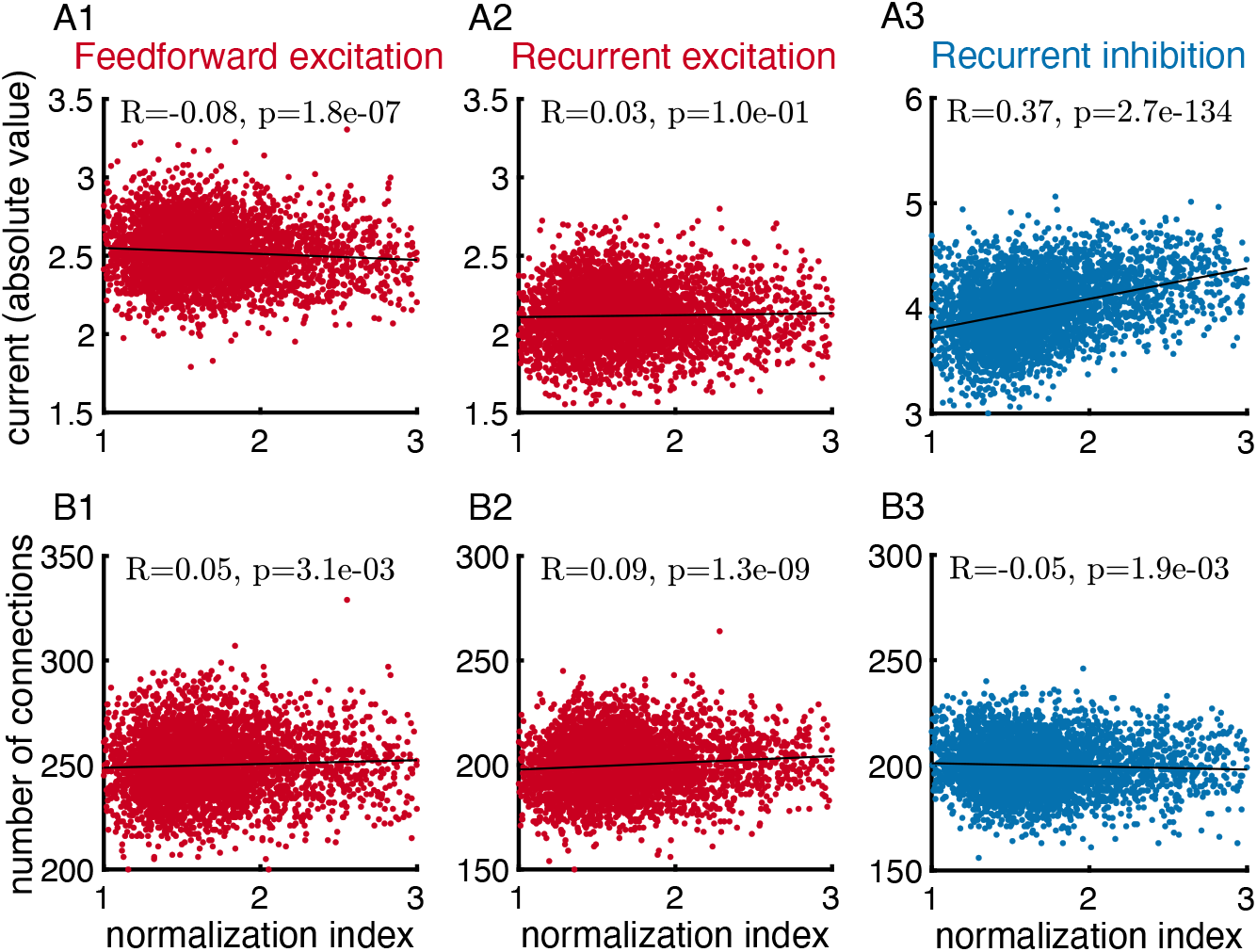
Related to Figure 3. The normalization index of neurons is strongly correlated with the magnitude of the mean recurrent inhibitory current. (**A**) There is a strong correlation between the firing rate normalization indexes and the average inhibitory current a neuron receives when two images are presented (**A3**), and only weak correlations with the feedforward (**A1**) and recurrent (**A2**) excitatory currents. (**B**) The correlation between normalization and the number of excitatory or inhibitory input connections is weak.

**Figure S8:**
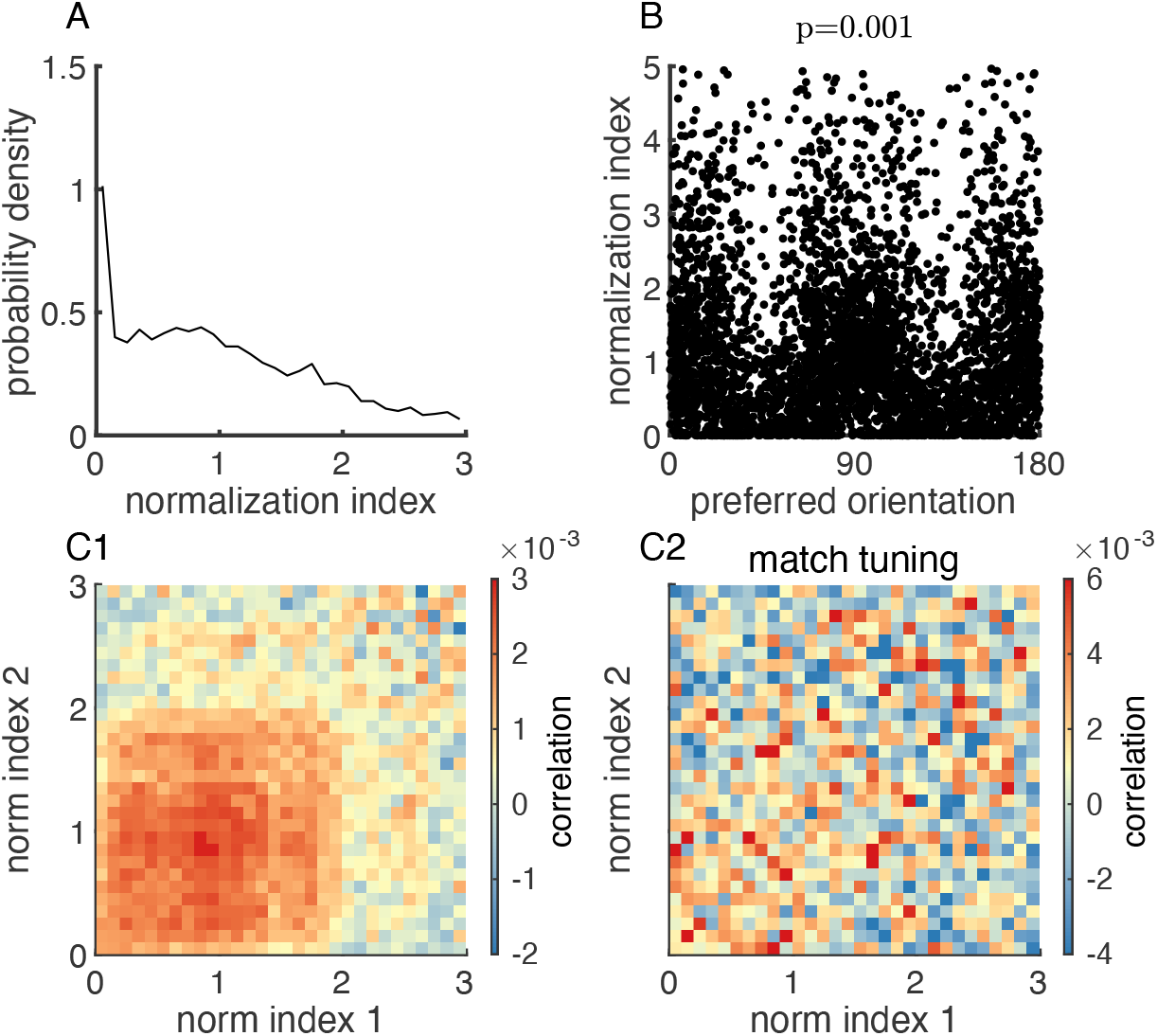
The normalization index of a neuron strongly depends on the neuron’s preferred orientation in the stabilized supralinear network (SSN) model. The SSN model is a large-scale, probabilistically connected, 2D model of a visual area. E/I units are arranged on a grid of 75 ×75. Preferred orientations are assigned according to a superposed orientation map. Parameters of the model are the same as those used in Figure 6 of Rubin et al., 2015. Additionally, we apply globally correlated additive noise to each unit, *ξ*_*i*_, where *ξ*_*i*_ satisfies 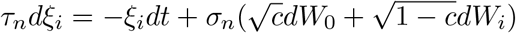, with *dW*_0_ and *dW*_*i*_ being Wiener process, *τ*_*n*_ = 40 ms, *σ*_*n*_ = 3.5, and correlation *c* = 0.2. (**A**) The normalization index of model neurons is broadly distributed in the SSN model. (**B**) The normalization index and the tuning preference of model SSN neurons are statistically dependent (*p* = 0.001). (**C1**) Spike count correlations between a pair of neurons as a function of the normalization indexes of the pair. (**C2**) The dependence of spike count correlation on normalization indexes is absent after matching for the distribution of tuning preferences of neurons across normalization indexes (**C2**).

